# A gene expression program activated by neuronal inactivity

**DOI:** 10.1101/636878

**Authors:** Zhonghua Zhu, Jennifer Lennon, Timothy M. Barry, Tamara Sanchez Ortiz, Seana Lymer, Shaul Mezan, Sebastian Kadener, Justin Blau

## Abstract

Long term synaptic plasticity requires transcriptional changes in response to neuronal activity. While neuronal activity rapidly increases transcription of activity-regulated genes, it is less clear how neurons respond to neuronal inactivity. The plasticity gene *Pura* encodes a Rho1 GEF whose rhythmic expression drives 24 hour rhythms in the retraction of the projections of *Drosophila* LN_v_ circadian pacemaker neurons. We found that *Pura* transcription in LN_v_s is repressed by neuronal activity and induced by neuronal inactivity – the opposite of canonical activity-regulated genes. We identified the Pax6 transcription factor Toy as the relevant activator of *Pura* transcription, and found that *toy* transcription itself is induced by neuronal inactivity and by reducing intracellular calcium. Thus we propose that neurons contain complementary gene expression programs that are activated by either increased or decreased neuronal activity.

## Introduction

Neuronal plasticity is essential for the nervous system to wire correctly and underlies processes including learning and memory and recovery from strokes. ^1,2^ Disruptions to plasticity are increasingly associated with neurological disorders such as autism spectrum disorder and schizophrenia.^3,4^

Plasticity includes changes in both the excitability and structure of neurons. While changes in neuronal excitability alter the strength of the signals transmitted, structural plasticity can create new neuronal connections. New gene expression plays a major role in plasticity via a set of activity regulated genes (also known as immediate early genes) whose transcription is rapidly induced by neuronal firing. The products of some activity regulated genes such as *Arc* and *Homer1a* directly regulate synaptic function, while others such as *c-fos*, *Nr4a1* and *Egr1* encode transcription factors whose targets likely include genes that alter neuronal connectivity.^5,6^

However, neuronal plasticity also includes reducing neuronal excitability and weakening or losing connections. These types of neuronal plasticity are probably important in updating memories and in sleep, in which synaptic downscaling likely reduces overall connectivity and synapse numbers.^7–9^ However, relatively few molecular mechanisms have so far been described to explain how neurons decrease connectivity, and the role of gene expression in this phenomenon has not been well-studied.

One example of how gene expression can weaken neuronal connections comes from *Drosophila* circadian pacemaker neurons, which drive ∼24hr rhythms in behavior.^10^ A subset of pacemaker neurons – the 4 small ventral lateral neurons (s-LN_v_s) in each brain hemisphere – show predictable and intrinsic daily rhythms in their excitability and in the structure of their projections.^11,12^. We found that Rho1 activity is rhythmic in s-LN_v_ projections, and this is regulated by circadian rhythms in the expression of *Puratrophin-like* (*Pura*), a Rho1 Guanine nucleotide exchange factor (GEF) that increases Rho1 activity.^13^

s-LN_v_ plasticity is also regulated by neuronal activity, which normally peaks around dawn^11^ when s-LN_v_ projections are maximally expanded. However, artificially inducing s-LN_v_s to fire at dusk rapidly expands their projections to the dawn-like state.^14^ Part of the mechanism that allows s-LN_v_ projections to expand at dusk in response to firing could be that increasing neuronal activity reduces *Pura* RNA levels in LN_v_s.^15,16^ Indeed, it seems important for neuronal activity to repress genes such as *Pura* that retract neuronal projections since activity-induced genes tend to have the opposite effect to *Pura* on neuronal structure.^6^

We investigated how *Pura* transcription responds to neuronal activity in intact brains and contrasted this with transcription of *Hr38*, a typical activity-regulated gene whose expression is induced by neuronal activity in invertebrates just like its mammalian orthologue *Nr4a1.*^17,18^ We confirmed that neuronal activity induced *Hr38* transcription in LN_v_ clock neurons. In contrast, neuronal activity repressed *Pura* transcription, while hyperpolarizing LN_v_s activated *Pura* transcription. Thus transcription of *Hr38* and *Pura* respond to neuronal activity and inactivity, but in opposite directions.

Dissecting the *Pura* regulatory region, we identified an 116bp enhancer that gives the correct spatial and temporal expression of *Pura* in LN_v_s and responds to both neuronal activity and hyperpolarization. This enhancer contains binding sites for Mef2, Stripe (Sr, the *Drosophila* Egr-1 orthologue) and Atf-3, which are either post-translationally or transcriptionally activated by neuronal activity in mammals.^19–21^ All three factors repressed *Pura* transcription, suggesting a mechanism for how neuronal activity represses *Pura*. We also identified that Toy, one of the two *Drosophila* Pax6 transcription factors^22^, activates *Pura* transcription likely through a binding site in the minimal *Pura* enhancer. Strikingly, *toy* transcription itself is activated when LN_v_s are hyperpolarized. Mechanistically, we show that reducing intracellular calcium is sufficient to activate *toy* transcription. Furthermore, *toy* is both necessary and sufficient to retract s-LNv projections. We speculate that inactivity-driven gene expression is a general property of plastic neurons.

## Results

### Transcription of *Hr38* and *Pura* responds in opposite directions to neuronal activity and hyperpolarization

We developed reporter genes to test if transcription of *Hr38* and *Pura* responds to neuronal activity. To identify the regulatory regions that direct expression to LN_v_s, we crossed available *Hr38-* and *Pura*-*Gal4* lines^23,24^ to *UAS-GFP* and identified one Gal4 line for each gene that expressed in larval LN_v_s (Fig. 1A and Fig. S1A). The *Hr38* enhancer from this Gal4 line was fused upstream of the *Hsp70* basal promoter, whereas the *Pura* enhancer already contained its endogenous basal promoter and transcriptional start site. Downstream we fused DNA encoding a tdTomato fluorescent protein that is codon-optimized for *Drosophila* and has a nuclear localization signal (NLS) to focus signals for quantification.^25^ This version of tdTomato was destabilized by adding a PEST sequence to report changes in transcription over time. We also added an intervening sequence (IVS) to promote nuclear export of spliced mRNA and a synthetic 5’ UTR (Syn21) to increase translation.^26^ ^27^ *Hr38-* and *Pura-Tomato* constructs were inserted to the same location as each other on chromosome 2.

**Figure 1.**
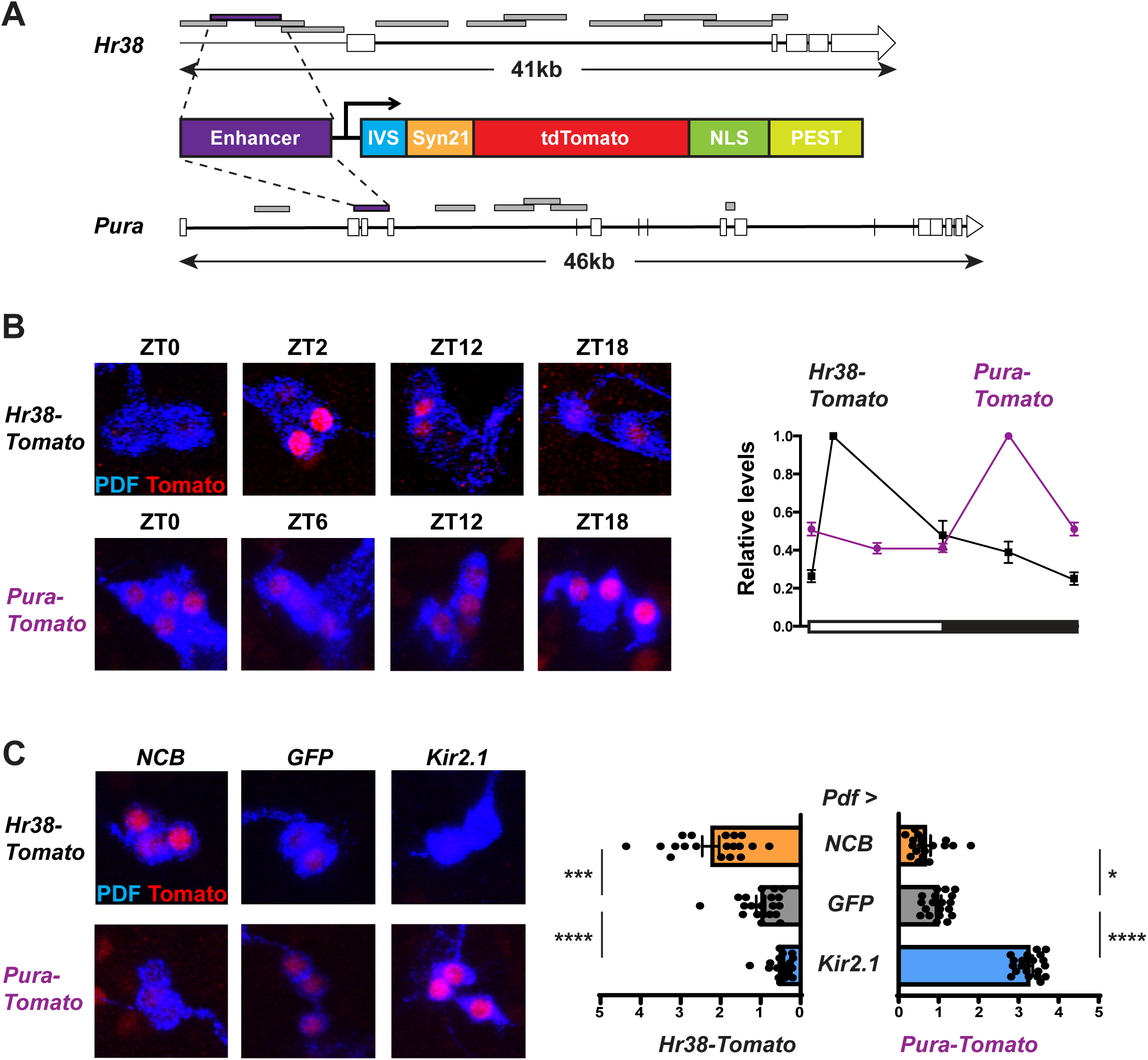
*Hr38* and *Pura* transcription respond in opposite directions to neuronal activity and hyperpolarization. **(A) Schematic of *Hr38* and *Pura* gene loci and reporter genes.** Gene structures of *Hr38* and *Pura* are shown with exons as white boxes, introns as thick black lines and intergenic regions as thin black lines. Enhancers tested are shown above the gene models and colored either grey (no expression in larval LN_v_s when crossed to GFP) or purple (expression in LN_v_s, Fig. S1A). For transcriptional reporter genes, enhancers were fused upstream of a basal promoter (transcription start site denoted by arrow), an intervening sequence (IVS, blue), a synthetic 5’UTR (Syn21, orange), tdTomato cDNA (red), 3 nuclear localization sequences (NLS, green) and a PEST sequence (yellow) to destabilize tdTomato.^25^ **(B) *Hr38* and *Pura* transcriptional reporters have differently phased rhythms.** Larvae with either *Hr38-* or *Pura-Tomato* transgenes were entrained in LD cycles at 25°C. Third instar larval brains were dissected at the times shown and stained with antibodies to Tomato (red) and to PDF to mark LN_v_s (blue). Left: Representative confocal images. The same confocal settings were applied to each scan for all timepoints in one experiment. Right: Tomato signals in a single LN_v_ were quantified and normalized to the average value at the peak time for each reporter. Error bars show SEM. The white:black bar on the x-axis represents time in 12:12 LD cycles. The data from ZT0 are replotted at ZT24. Levels of *Hr38-* (black line) *and Pura-Tomato* (purple line) vary with time in larval LN_v_s (one-way ANOVA, p < 0.0001). The graph is an average of 2 independent experiments with 15-20 single LN_v_s from 5 brains in each experiment. **(C) *Hr38* and *Pura* transcriptional reporter genes respond to long-term hyperexcitation and hyperpolarization in opposite directions.** Tomato levels were measured in LN_v_s at ZT24 from larvae with either *Hr38-* or *Pura-Tomato* transgenes and *Pdf-Gal4* (*Pdf >*) expressing *UAS-NaChBac* (*NCB*) for hyperexcitation, *UAS-GFP* (*GFP*) as control or *UAS-Kir2.1* (*Kir2.1*) for hyperpolarization. Larval brains were dissected, stained and quantified as in Fig. 1B with representative confocal images on the left and quantification on the right. Tomato signals in single LN_v_s were normalized to the average of the control group. *Hr38-Tomato* levels are higher in *NCB*-expressing LN_v_s (orange bars) and lower in *Kir2.1*-expressing LN_v_s (blue bars) than in *GFP*-expressing LN_v_s (grey bars). Student’s *t* test: *NCB* vs *GFP* p < 0.0001, *Kir2.1* vs *GFP* p < 0.001). *Pura-Tomato* levels are lower in *NCB*-expressing LN_v_s and higher in *Kir2.1*-expressing LN_v_s than in *GFP*-expressing LN_v_s. Student’s *t*-test: *NCB* vs *GFP* p < 0.05, *Kir2.1* vs *GFP* p < 0.0001). Error bars show SEM. The graph is an average of 2 independent experiments with 15-20 single cells from 5 brains in each experiment.

We first measured Tomato levels in larval LN_v_s across a 12hr:12hr light:dark (LD) cycle. We found that *Hr38-Tomato* levels in LN_v_s peak at ZT2 (Zeitgeber Time, ZT0 = lights on, ZT12 = lights off) (Fig. 1B). This is consistent with previous reports of higher *Hr38* mRNA levels in LN_v_s at CT3 than CT15 (Circadian Time, time in constant darkness after entrainment in LD cycles)^15^ and with LN_v_s being more excitable at dawn than dusk.^11^ *Pura-Tomato* levels reached a maximum at ZT18 and were low by ZT24 (Fig. 1B), which is similar to *Pura* RNA.^28^ The data in Fig.1B are for *Pura-Tomato B*, which has ∼1.5kb of regulatory region directly upstream of the *Pura* basal promoter. *Pura-Tomato B* showed very similar temporal dynamics as *Pura-Tomato A*, which has an additional ∼0.5kb downstream of the transcriptional start site (Fig. S1B). Thus *Hr38* and *Pura* transcriptional reporters have similar dynamics to their RNAs.^15,28^

To test how neuronal activity regulates *Hr38* and *Pura* transcription, we first used a *UAS-NaChBac* transgene (*UAS-NCB*) to hyperexcite LN_v_s. *NCB* is a low threshold voltage-gated bacterial Na^+^ channel that increases the excitability of LN_v_s.^29,30^ *UAS-NCB* was expressed specifically in larval LN_v_s using the *Pigment dispersing factor-Gal4* driver (*Pdf-Gal4*) ^31^. We use the nomenclature *Pdf > X* to indicate *Pdf-Gal4* and *UAS-X* transgenes in the figures. Larvae were entrained in LD and *Hr38-Tomato* and *Pura-Tomato* levels were measured at ZT24, just before the normal increase in *Hr38* transcription shortly after dawn. Expressing NCB in LN_v_s increased *Hr38-Tomato* levels and modestly reduced *Pura-Tomato* levels compared to control LN_v_s expressing a *UAS-GFP* transgene (Fig. 1C).

To test how hyperpolarization affects *Hr38* and *Pura* transcription, we used *Pdf-Gal4* to express *UAS-Kir2.1*, a mammalian inward rectifier K^+^ channel.^32^ We again measured *Hr38-* and *Pura-Tomato* levels at ZT24 and found that *Hr38-Tomato* levels were lower than in control LN_v_s, while *Pura-Tomato* levels were higher in Kir2.1-expressing LN_v_s than in control LN_v_s (Fig. 1C). Thus *Hr38* and *Pura* respond in opposite directions to long-term changes in neuronal activity: *Hr38* transcription is activated when LN_v_s are hyperexcited and repressed when LN_v_s are hyperpolarized, whereas *Pura* is activated by hyperpolarization and repressed by increased activity.

### *Hr38* and *Pura* transcription responds in opposite directions to transient changes in neuronal activity

To test if *Hr38* and *Pura* transcription respond to more transient changes in neuronal activity, we used *Pdf-Gal4* to ectopically express *UAS-dTrpA1*, a heat-activated *Drosophila* cation channel that is inactive below 23°C.^33^ Larvae expressing TrpA1 in LN_v_s were entrained in LD cycles at 22**°**C and then shifted to 30**°**C for 4hr from ZT12 to ZT16. We selected lights-off to induce firing since LN_v_s are relatively inactive at this time.^11^ The data in Fig. 2A show that *Hr38-Tomato* levels increased in *dTrpA1*-expressing LN_v_s compared to control LN_v_s without a *dTrpA1* transgene that followed the same temperature shift. In contrast, *Pura-Tomato* levels were repressed by dTrpA1-induced neuronal activity.

**Figure 2.**
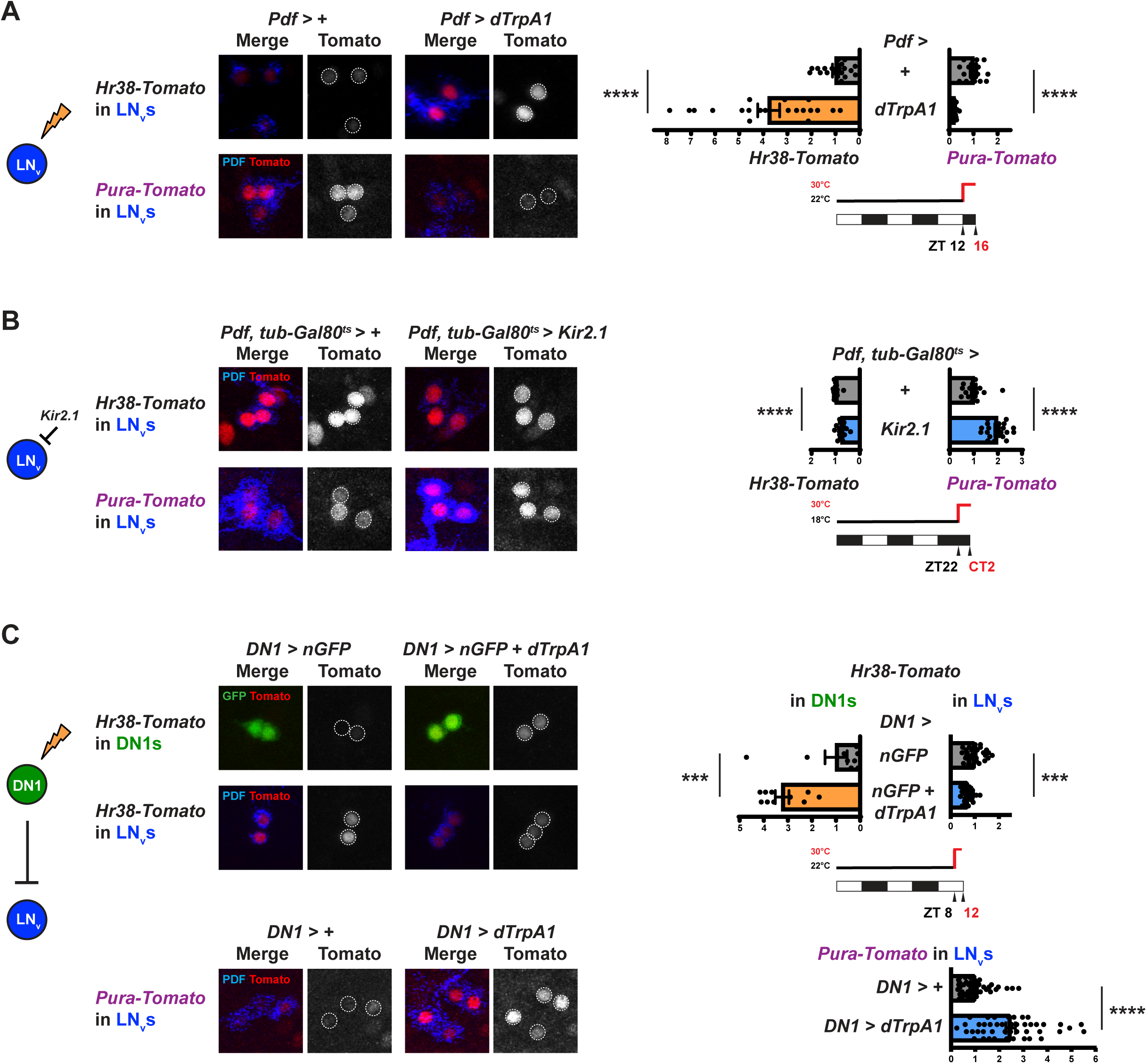
*Hr38* and *Pura* transcription respond in opposite directions to transient changes in neuronal activity. **(A) Transcriptional responses to inducing LN_v_ firing.** Larvae with *Pdf-Gal4* and either *Hr38-*or *Pura-Tomato* were raised at 22°C in LD cycles. Control larvae had no *UAS* transgene (*Pdf > +*). Experimental larvae had a *UAS-dTrpA1* transgene (*Pdf > dTrpA1*). Larvae were shifted to 30°C from ZT12 to ZT16 to activate dTrpA1 and induce LN_v_ firing – see timeline on bottom right. Larval brains were dissected at ZT16 and stained and quantified as in Fig. 1B. Left: representative confocal images with Tomato (red) and PDF to mark LN_v_s (blue) on left and Tomato single channel images on the right. Dashed lines circle LN_v_ nuclei for quantification. Right: quantification of 2 independent experiments with 15-20 single cells from 5 brains in each experiment*. Hr38-Tomato* levels are higher in *dTrpA1*-expressing LN_v_s (orange bars) than in control LN_v_s (grey bars). Student’s *t*-test, p < 0.0001). *Pura-Tomato* levels are lower in *dTrpA1*-expressing LN_v_s than in control LN_v_s (Student’s *t*-test, p < 0.0001). Error bars show SEM. **(B) Responses to inducing LN_v_ hyperpolarization.** Larvae with *Pdf-Gal4*, *tub-Gal80^ts^* and either *Hr38-* or *Pura-Tomato* were raised at 18°C in LD cycles. Control larvae had no *UAS* transgene (*Pdf, tub-Gal80^ts^ > +*). Experimental larvae had a *UAS-Kir2.1* transgene whose expression was suppressed at 18°C by Gal80 (*Pdf, tub-Gal80^ts^ > Kir2.1*). Larvae were shifted to 30°C at ZT22 and maintained in the dark for 4hr before dissection at CT2 – see timeline, bottom right. Staining and quantification were as in Fig. 1B. Left: representative confocal images as in Fig. 2A. Right: quantification of 2 independent experiments with 15-20 single cells from 5 brains in each experiment. *Hr38-Tomato* levels were lower in *Kir2.1*-expressing LN_v_s (blue bars) than in control LN_v_s (grey bars; Student’s *t*-test, p < 0.0001). *Pura-Tomato* levels were higher in *Kir2.1*-expressing LN_v_s than in control LN_v_s (Student’s *t*-test, p < 0.0001). Error bars show SEM. **(C) Transcriptional responses in LN_v_s and DN1s to activating DN1s.** Larvae with *cry39-Gal4* and *Pdf-Gal80* expressing UAS transgenes in DN1s (*DN1 >*) and either *Hr38-Tomato or Pura-Tomato* reporter genes were raised at 22°C in LD cycles. Larvae were shifted to 30°C at ZT8 for 4hr before dissection at ZT12 – see timeline. Larval brains were stained with antibodies to GFP to mark DN1s (green), PDF to mark LN_v_s (blue) and Tomato (red). Quantification as in Fig. 1B with averages from 2 independent experiments containing 10 DN1s or 15-20 LN_v_s from 5 brains in each experiment. Far left: illustration shows that inducing neuronal activity in DN1s (green) inhibits LN_v_ (blue). Left: Representative confocal images are shown as in Fig. 2A. Top left: Control larvae had *UAS-nGFP* (*DN1 > nGFP*) and the experimental group had *UAS-nGFP* and *UAS-dTrpA1* transgenes (*DN1 > nGFP* + *dTrpA1*). *Hr38-Tomato* levels were higher in *dTrpA1*-expressing DN1s than in control DN1s (Student’s *t*-test, p < 0.001) but lower in LN_v_s in which DN1s were induced to fire (Student’s *t*-test, p < 0.001). Bottom left: Genotypes as above except that the reporter gene was *Pura-Tomato*. Levels of *Pura-Tomato* were higher in LN_v_s of larvae with dTrpA1 activated in DN1s than in control LN_v_s (Student’s *t*-test, p < 0.0001). Error bars show SEM.

We also tested the responses of *Hr38* and *Pura* to transient hyperpolarization. For this, we used a temperature-sensitive Gal80 transgene (*tubulin-Gal80^ts^* ^34^) to repress Gal4 activity and control the timing of expression of *UAS-Kir2.1*. Larvae were entrained in LD cycles at 18**°**C when Gal80^TS^ is active, and then shifted to 30**°**C at ZT22 to inactivate Gal80 and induce *Kir2.1* expression. Larvae were maintained at 30**°**C for 4hr in darkness to prevent light-induced neuronal activity.^30^ The data in Fig. 2B show that inducing *Kir2.1* expression in LN_v_s decreased *Hr38-Tomato* levels relative to control LN_v_s. In contrast, *Pura-Tomato* levels were higher in LN_v_s expressing *Kir2.1* than in control LN_v_s. Thus *Hr38* and *Pura* transcription respond in the same direction to more transient changes (Fig. 2A, B) as they do to long-term changes in LN_v_ electrical activity (Fig. 1C).

### *Hr38* and *Pura* transcription respond in opposite directions to hyperpolarization via an intact neural circuit

The DN1 dorsal larval clock neurons release glutamate and inhibit LN_v_s via GluCl, a glutamate-gated chloride channel.^35^ We decided to increase DN1 activity and ask if changes in *Hr38* and *Pura* transcription in LN_v_s could be detected when LN_v_ activity was inhibited via a neural circuit. We used the combination of *cry39-Gal4* and a temperature-independent *Pdf-Gal80* transgene to restrict transgene expression to larval DN1s. *cry39-Gal4* is expressed in larval DN1s and LN_v_s^36^, but the *Pdf-Gal80* transgene constitutively blocks Gal4 activity in LN_v_s.^37^ We confirmed that this driver combination represses *UAS-GFP* transgene expression in LN_v_s (Fig. S2). Larvae with *cry39-Gal4*, *Pdf-Gal80* and *UAS-dTrpA1* transgenes are labeled in Fig. 2C as *DN1 > TrpA1* for simplicity.

Larvae were entrained in LD cycles at 22**°**C and then shifted to 30**°**C for 4hr from ZT8 to ZT12 to activate dTrpA1 in DN1s. *Hr38-Tomato* also expresses in DN1s. The data in Fig. 2C show that *Hr38-Tomato* levels at ZT12 were higher in *dTrpA1*-expressing DN1s than in control DN1s that went through the same temperature shift but lacked a *dTrpA1* transgene. These data confirm that dTrpA1 induced DN1 firing. We also measured *Hr38-Tomato* levels in LN_v_s. The data in Fig. 2C show that *Hr38-Tomato* levels were lower in LN_v_s in larvae in which DN1s were induced to fire than in LN_v_s from control larvae (Fig. 2C). Together, these data are consistent with DN1s inhibiting LN_v_ activity.

Finally, we measured *Pura-Tomato* levels in LN_v_s when DN1s were induced to fire from ZT8 to ZT12. The data in Fig. 2C show that *Pura-Tomato* levels in LN_v_s at ZT12 were higher in larvae with active dTrpA1 in DN1s than in control larvae without a *dTrpA1* transgene. Thus inhibiting LN_v_ activity by activating DN1s increases *Pura* transcription in LN_v_s. These data support the idea that *Pura* transcription can be activated by transiently decreasing neuronal activity, and show that this can occur in the context of an intact neural circuit.

### Dissecting the *Pura* enhancer to identify cis-regulatory elements

To determine how neuronal inactivity activates *Pura* transcription, we first sought to find the minimal *Pura* enhancer that responds to changes in neuronal activity in LN_v_s to identify the relevant cis-regulatory elements. The 1.5kb *Pura-B* enhancer in Fig. 1B contains the likely transcription start site for *Pura* transcript B and our deletion constructs all used 100bp surrounding this transcription start site as a basal promoter (shown in yellow in Fig. 3A) with varying lengths of upstream DNA. The different constructs were inserted into the same chromosomal location as *Pura-Tomato B* and assayed in larval LN_v_s.

**Figure 3.**
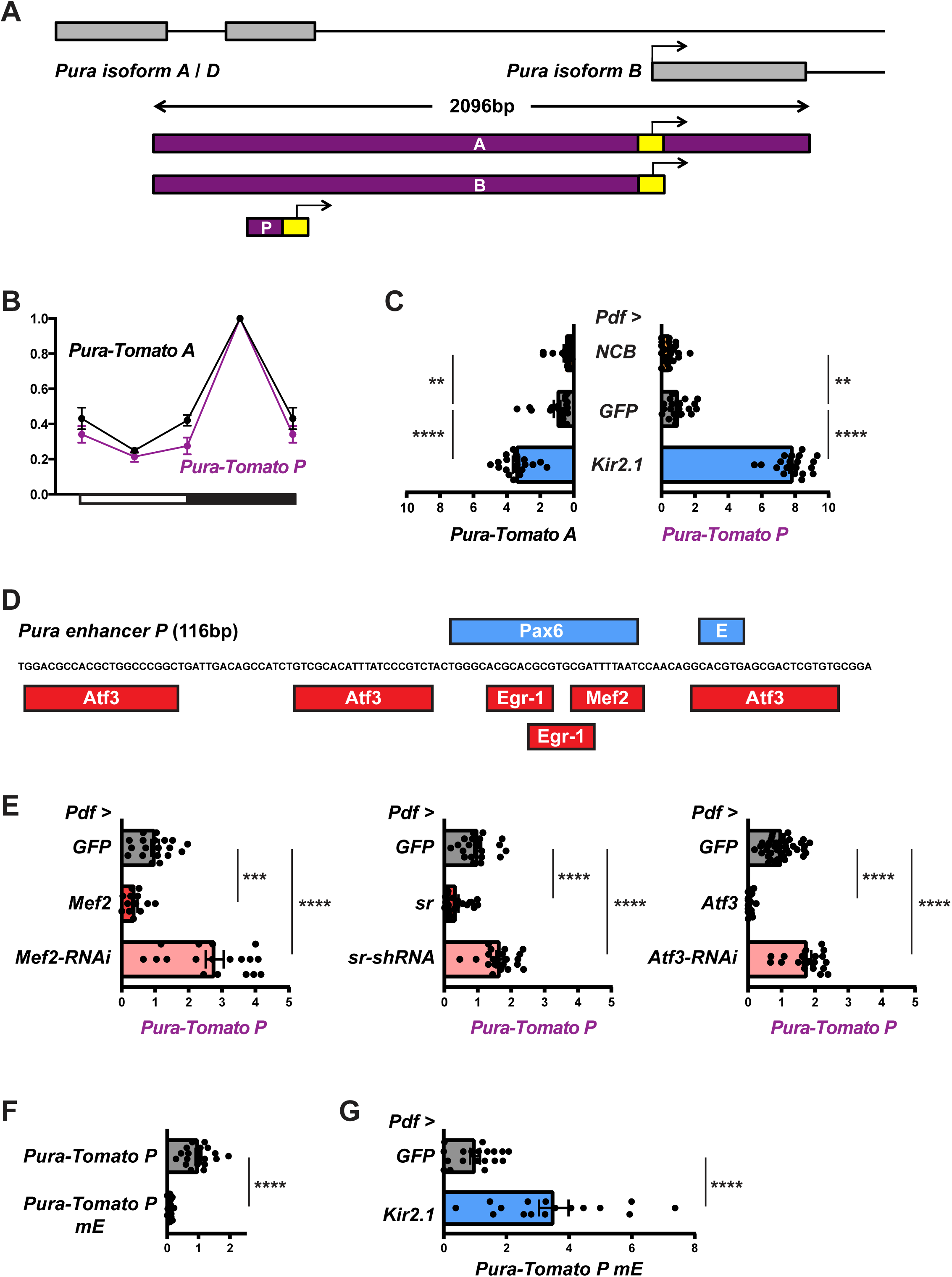
The *Pura* enhancer contains cis-regulatory elements for activity-regulated transcription factors. **(A) A minimal *Pura* enhancer for LN_v_ expression.** Map of the *Pura* regulatory region showing exons (grey boxes), introns (black lines) and the transcription start site (TSS) marked with an arrow. *Pura* enhancers (purple) were fused with the *Pura* basal promoter (−50bp to +50bp of TSS, yellow) to drive Tomato expression. This map shows the enhancer used in the original 2kb *Pura-Tomato A* reporter, the 1.5kb *Pura-Tomato B* enhancer that lacks the 5’UTR, and the 116bp enhancer in *Pura-Tomato P*. **(B) *Pura-Tomato A* and *Pura-Tomato P* have similar phase rhythms in larval LN_v_s.** Transgenic larvae with either *Pura-Tomato A* (black line in graph) or *Pura-Tomato P* (purple line) were entrained in LD cycles at 25°C and brains dissected at the times shown. Staining, quantification and normalization are as in Fig. 1B with data from ZT0 replotted at ZT24. *Pura-Tomato A* and *P* levels vary with time in LN_v_s (one-way ANOVA, p < 0.0001). Error bars show SEM. The graph is an average of 2 independent experiments with 15-20 LN_v_s from 5 brains in each experiment. **(C) *Pura-Tomato A* and *Pura-Tomato P* respond similarly to neuronal activity.** Larvae with *Pdf-Gal4* expressing *UAS-NaChBac* (*NCB*), *UAS-GFP* (*GFP*, black) or *UAS-Kir2.1* (*Kir2.1*, blue), and either *Pura-Tomato A* or *Pura-Tomato P* were entrained in LD cycles. Larval brains were dissected at ZT12, stained, quantified and the data normalized as in Fig. 1B. Left: *Pura-Tomato A* levels are lower in *NCB*-expressing LN_v_s than in *GFP*-expressing LN_v_s. *Pura-Tomato A* levels are higher in *Kir2.1*-expressing LN_v_s than in *GFP*-expressing LN_v_s. Student’s *t*-tests: *NCB* vs *GFP* p < 0.05, *Kir2.1* vs *GFP* p < 0.0001. Right: *Pura-Tomato P* levels are lower in *NCB*-expressing LN_v_s than in *GFP*-expressing LN_v_s. *Pura-Tomato P* levels are higher in *Kir2.1*-expressing LN_v_s than in *GFP*-expressing LN_v_s. Student’s *t*-tests, *NCB* vs *GFP* p < 0.01, *Kir2.1* vs *GFP* p < 0.0001. Error bars show SEM. The graph shows an average of 2 independent experiments with 15-20 LN_v_s from 5 brains in each experiment. **(D) cis-regulatory elements in *Pura enhancer P*.** The DNA sequence of the 116bp *Pura enhancer P* along with predicted binding sites for potential activators (Pax6 and E for E-box) in blue above the DNA sequence and potential repressors (Atf3, Egr-1 and Mef2) in red below the DNA sequence. **(E) *Pura-Tomato P* is repressed by Mef2, Sr and Atf3.** *Pura-Tomato P* levels were measured in LN_v_s at ZT12 from larvae with *Pdf-Gal4* expressing *UAS-Dcr-2* and a UAS transgene to either overexpress Mef2, Sr or Atf3 (red bars) or an RNAi transgene to reduce expression (pink bars). *UAS-GFP* (*GFP*) was used as a control (grey bars). Larval brains were dissected, stained, quantified and the data normalized as in Fig. 1B. Left: *Pura-Tomato P* levels are lower in LN_v_s over-expressing *Mef2* (*Pdf > Mef2*) and higher in *Mef2-RNAi*-expressing LN_v_s (*Pdf > Mef2-RNAi*) than in control LN_v_s (*Pdf > GFP*). Student’s *t*-test: *Pdf > Mef2* vs *Pdf > GFP*, p < 0.001; *Pdf > Mef2-RNAi* vs *Pdf > GFP*, p < 0.0001. Middle: *Pura-Tomato P* levels are lower in LN_v_s over-expressing *sr* (*Pdf > sr*) and higher in *sr-shRNA*-expressing LN_v_s (*Pdf > sr-shRNA*) than in control LN_v_s (*Pdf > GFP*). Student’s *t*-test: *Pdf > sr* vs *Pdf > GFP* and *Pdf > sr-shRNA* vs *Pdf > GFP*, p < 0.0001. Right: *Pura-Tomato P* levels are lower in LN_v_s over-expressing *Atf3* (*Pdf > Atf3*) and higher in *Atf3-RNAi*-expressing LN_v_s (*Pdf > Atf3-RNAi*) than in control LN_v_s (*Pdf > GFP*). Student’s *t*-test: *Pdf > Atf3* vs *Pdf > GFP* and *Pdf > Atf3-RNAi* vs *Pdf > GFP*, p < 0.0001. Error bars show SEM. The graphs show an average of 2 independent experiments with 15-20 LN_v_s from 5 brains in each experiment. **(F) An E-box in *Pura enhancer P* contributes to expression in LN_v_s.** Larvae with either wild type *Pura-Tomato P* or with the core of the E-box mutated (*Pura-Tomato P mE*) were entrained in LD cycles at 25°C. Larval brains were dissected at ZT12 and stained, quantified and the data normalized as in Fig. 1B. *Pura-Tomato P* levels in LN_v_s are higher than *Pura-Tomato P mE* (Student’s *t*-test, p < 0.0001). The graph is an average of 2 independent experiments with 15-20 LN_v_s from 5 brains in each experiment. **(G) The E-box in the minimal *Pura* enhancer is not required for transcriptional activation by Kir2.1.** *Pura-Tomato P mE* levels were measured in LN_v_s at ZT12 from larvae with *Pdf-Gal4* expressing either *UAS-GFP* (*Pdf > GFP*, grey bar) or *UAS-Kir2.1* (*Pdf > Kir2.1*, blue bar). Larval brains were dissected, stained, quantified and the data normalized as in Fig. 1B. *Pura-Tomato P mE* levels are higher in *Kir2.1*-expressing LN_v_s than in control LN_v_s (Student’s *t*-test, p < 0.0001). Error bars show SEM. The graph is an average of 2 independent experiments with 15-20 LN_v_s from 5 brains in each experiment.

To identify the minimal *Pura* enhancer that retains the spatial and temporal expression pattern in LN_v_s, larvae were entrained in LD and *Pura-Tomato* levels measured at ZT18, the time of peak expression (Fig. 1B). Fig. 3A and Fig. S3A show a series of deletions of the *Pura* enhancer with purple indicating expression and blue denoting no detectable expression in LN_v_s. *Pura-Tomato P* was the smallest transgene that expressed in LN_v_s and contains only 116bp upstream of regulatory DNA upstream of the basal promoter. *Pura-Tomato P* has the same temporal pattern in LN_v_s as *Pura-Tomato A* (Fig. 3B), as do the other deletions of the regulatory regions that express in LN_v_s (Fig. S3B).

We then tested if *Pura-Tomato P* responds to neuronal activity and hyperpolarization like the larger constructs. We used *Pdf-Gal4* to express *UAS-GFP* (as a control), *UAS-NCB* or *UAS-Kir2.1* and measured *Pura-Tomato A* and *Pura-Tomato P* levels in LN_v_s at ZT12. The data in Fig. 3C show that *Pura-Tomato P* was induced by expressing Kir2.1 and repressed by NCB just like *Pura-Tomato A*. Thus we focused on *Pura-Tomato P* to identify cis-regulatory elements that respond to neuronal activity and neuronal inactivity.

### *Pura* is repressed by activity-regulated transcription factors

The 116bp *Pura* enhancer is highly conserved across 23 *Drosophila* species^38^, although this is partly because this region overlaps with a protein coding exon from *Pura* RNA isoforms A and D (Fig. 3A). To predict cis-regulatory elements in the *Pura* enhancer *P*, we used MATCH analysis from TRANSFAC ^39^ and focused on transcription factors known to be regulated by neuronal activity. This identified one potential Mef2 binding site, two potential Egr-1 binding sites and three potential Atf3 binding sites in *Pura P* (Fig. 3D). These sites are conserved in *D. pseudoobscura* and *D. virilis* (Fig. S3C).

While Mef2 is post-translationally modified by neuronal activity^40^, expression of *Egr-1* and *Atf3* are rapidly induced by neuronal activity in mammals.^20,21^ Expression of the fly *Egr-1* homolog – *stripe* (*sr*) – is rapidly induced by neuronal activity in adult LN_v_s^16^ and *sr* mRNA is higher when larval LN_v_s are more excitable at CT3 than at CT15.^15^ Similarly, *Atf3* mRNA levels in LN_v_s are also higher at CT3 than CT15.^15^

We performed gain-and loss-of-function experiments to test if these transcription factors regulate the minimal *Pura* enhancer. We measured *Pura-Tomato P* levels at ZT12 using *Pdf-Gal4* to either overexpress these transcription factors in LN_v_s or to express RNAi transgenes. The data in Fig. 3E show that *Pura-Tomato P* levels in LN_v_s were reduced by over-expressing Mef2, Sr or Atf3. Over-expressing Atf3 gave the strongest repression, consistent with the higher number of potential Atf3 binding sites (Fig. 3D). We also found that *Pura-Tomato P* levels increased when RNAi targeting any one of these transcription factors was expressed in LN_v_s (Fig. 3E). Thus we conclude that Mef2, Sr and Atf3 all repress *Pura* transcription.

### An E-box in the *Pura* enhancer is not essential for transcriptional activation by hyperpolarization

One sequence that stood out as a potential binding site for a transcriptional activator in the minimal *Pura* enhancer was a CACGTG E-box. This sequence can be directly bound by a heterodimer of Clock (Clk) and Cycle (Cyc), two bHLH transcription factors that are core clock components. Clk and Cyc drive circadian rhythms in the expression of many genes by interacting with the rhythmically produced repressor protein Period that regulates Clk/Cyc DNA-binding.^41^

To test the importance of this E-box in the response to hyperpolarization, we mutated its core to CA**GC**TG as in ref.^42^ and called this reporter gene *Pura-Tomato P mE* for **m**utant E-box. The data in Fig. 3F show that *Pura-Tomato P mE* levels were lower than *Pura-Tomato P* levels in LN_v_s at ZT12. To test whether the E-box is necessary for the response to hyperpolarization, we used *Pdf-Gal4* to constitutively express *Kir2.1* and compared levels of *Pura-Tomato P mE* at ZT12 with control LN_v_s expressing a GFP transgene. We found that levels of *Pura-Tomato P mE* were higher in *Kir2.1*-expressing LN_v_s than in control LN_v_s (Fig. 3G). Thus the E-box in the minimal *Pura* enhancer affects *Pura* expression, but is not necessary for transcriptional activation by hyperpolarization. We did not test if increasing neuronal activity represses *Pura-Tomato P mE* because expression of this reporter gene was too low in control LN_v_s to reliably measure potential decreases.

### *Pura* is activated by the Pax6 transcription factor Toy

As a complementary approach to identifying how *Pura* is regulated, we also performed an RNAi screen in LN_v_s. We focused on a set of 110 transcription factors that are expressed in fly heads ^43–45^ and for which there were available short hairpin RNA (shRNA) lines in a common genetic background ^46^. We used *Pdf-Gal4* to express *shRNA* transgenes against each of these transcription factors, and measured *Pura-Tomato B* reporter levels in LN_v_s at its peak at ZT18. Only three lines changed *Pura-Tomato B* levels by more than two standard deviations from the mean of control LN_v_s (Fig. 4A, Table S1). Two of these lines increased *Pura-Tomato B* levels modestly – these lines targeted *trachealess* and *pointed*. The third line stood out as it was the only line to reduce *Pura-Tomato B* expression, and also because it affected reporter gene levels so dramatically. This shRNA targets *twin-of-eyeless* (*toy*), and it reduced Tomato levels 5.5-fold compared to control LN_v_s, with the next strongest reduction of *Pura-Tomato B* only 1.4-fold (Fig. 4A).

**Figure 4.**
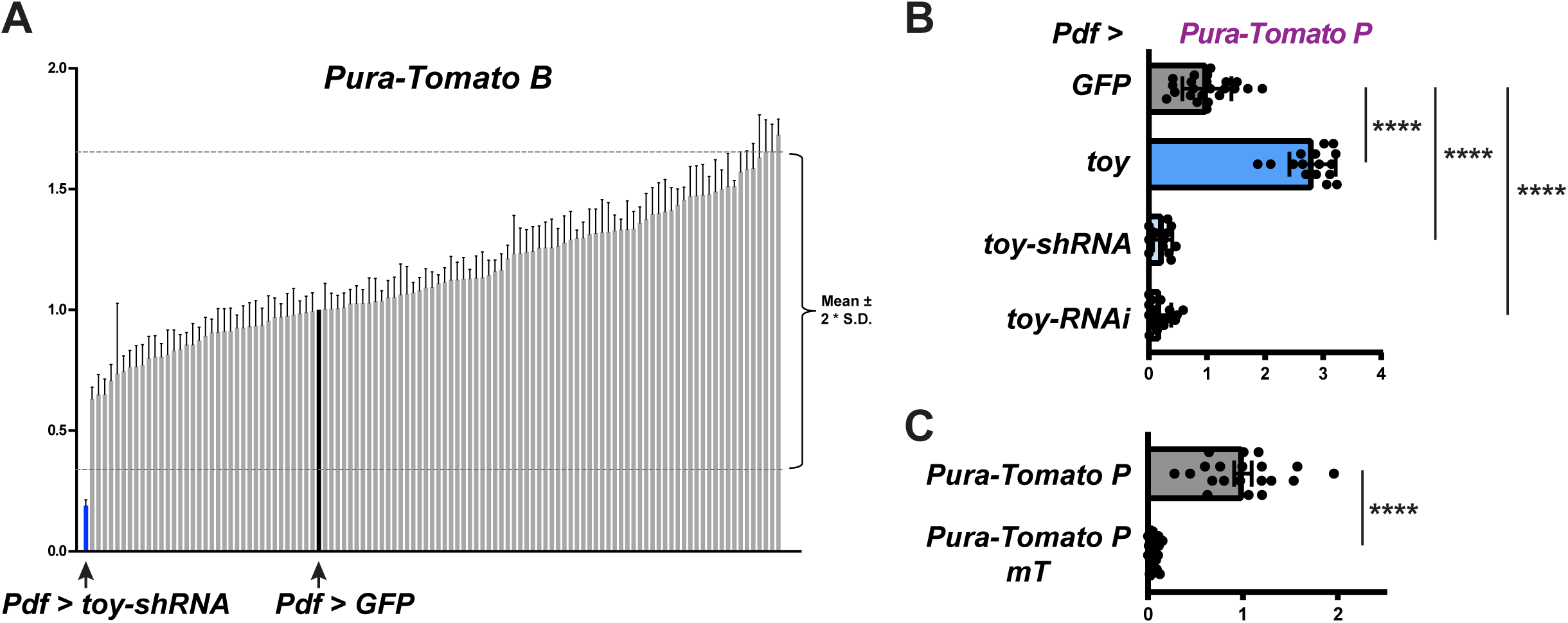
Toy is a transcriptional activator of *Pura*. **(A) An RNAi screen identified the transcription factor Toy as an activator of *Pura*.** Larvae with *Pura-Tomato B*, *Pdf-Gal4*, *UAS-Dcr-2* and a control transgene *UAS-GFP* (*GFP*) or one of 110 *UAS-shRNA* transgenes directed against a set of transcription factors were entrained in LD cycles at 25°C. Larval brains were dissected at ZT18 and stained and quantified as in Fig. 1B. Tomato signals in single LN_v_s were normalized to the mean of the *GFP*-expressing control LN_v_s (black line). Each grey bar represents the average *Pura-Tomato B* levels in 10-16 LN_v_s from 4 brains with one *UAS-shRNA* transgene. Error bars show SEM. Dashed lines show the mean of *Pura-Tomato B* levels in control LN_v_s ± 2 S.D. The lowest levels of *Pura-Tomato B* were in *UAS-toy*-*shRNA* expressing LN_v_s (blue line). Student’s *t*-test, p < 0.0001 vs control LN_v_s. **(B) The minimal *Pura* enhancer is activated by Toy.** *Pura-Tomato P* levels were measured at ZT12 in larvae with *Pdf-Gal4* expressing *UAS-Dcr-2* and a control *UAS-GFP* transgene (*Pdf > GFP*, grey bar), overexpressing *toy* via *UAS-toy* (*Pdf > toy*, blue bar), or reducing *toy* expression via *UAS-toy-shRNA* or *UAS-toy-RNAi* transgenes (*Pdf > toy-shRNA* and *Pdf > toy-RNAi*, pale blue bars). Larval brains were dissected, stained, quantified and the data normalized as in Fig. 1B. *Pura-Tomato P* levels are higher in *Pdf > toy* LN_v_s and lower with either *toy-shRNA* or *toy-RNAi* expressed in control LN_v_s. Student’s *t*-tests: p < 0.0001 for *Pdf > toy* vs *Pdf > GFP, Pdf > toy-shRNA* vs *Pdf > GFP* and *Pdf > toy-RNAi* vs *Pdf > GFP*. Error bars show SEM. The graph is an average of 2 independent experiments with 15-20 LN_v_s from 5 brains in each experiment. **(C)** The predicted Toy binding site in *Pura P* is required for expression in LN_v_s. Larvae with *Pura-Tomato P* or with the Toy binding site mutated (*Pura-Tomato P mT*) were entrained in LD cycles. Larval brains were dissected at ZT12 and stained as in Fig. 1B. Tomato signals in single LN_v_s were quantified and normalized to the average of the signals from *Pura-Tomato P* LN_v_s. *Pura-Tomato P* levels in LN_v_s are higher than *Pura-Tomato P mT* (Student’s *t*-test, p < 0.0001). The graph shows an average of 2 independent experiments with 15-20 LN_v_s from 5 brains in each experiment.

*toy* encodes one of the two *Drosophila* Pax-6 transcription factors. These proteins have two separate DNA binding domains: a Paired domain and a Homeodomain.^47^ *toy* is best known as a master regulator of eye development^22^ and it is also involved in mushroom body development.^48^ However, *toy* is also expressed in various neurons in adult flies, including LN_v_s^49^ and mushroom body neurons^50^, both of which are sites of plasticity.^12,51^

To test if Toy regulates the minimal 116bp *Pura* enhancer, we used *Pdf-Gal4* to express *UAS-toy* for overexpression, or *UAS-toy-shRNA* and *UAS-toy-RNAi* transgenes to reduce *toy* expression. *toy-shRNA* and *toy-RNAi* target different regions of *toy*. The results in Fig. 4B show that *Pura-Tomato P* levels at ZT12 in LN_v_s were higher in *toy*-overexpressing LN_v_s, and lower with *toy-shRNA* or *toy-RNAi*. Thus we conclude that Toy activates *Pura* transcription.

We searched for Pax-6 binding sites in the minimal *Pura* enhancer and found one Pax-6 binding site that overlaps with two of the Egr-1 sites and the Mef2 site (Fig. 3D). To test its function, we mutated this Pax-6 site in the context of *Pura-Tomato P*. This mutation also affects the predicted Egr-1 and Mef2 binding sites. Levels of the resulting construct (*Pura-Tomato P mT*) were almost undetectable in LN_v_s even at the normal peak of *Pura* expression at ZT18 (Fig. 4C). Thus this Pax-6 site is required for *Pura* expression and leads us to propose that Toy directly regulates *Pura* expression. *Pura-Tomato P mT* could not be activated with *Pdf-Gal4* expressing *UAS-Kir2.1* in LN_v_s (Fig. S4). However, the very low levels of expression of the *Pura-Tomato P mT* reporter gene prevent us making a strong conclusion that this Pax-6 site in *Pura* is required for induction by neuronal inactivity.

### *toy* transcription is activated by neuronal inactivity

Since the repressors that regulate *Pura* expression are likely regulated by post-translational modification (Mef2) or gene expression (Sr, Atf3), we wanted to test if *toy* is also regulated in LNvs. We first measured Toy protein levels in LN_v_s at different times of day and found that Toy levels change over 24hr in LN_v_s (Fig. 5A), peaking at ZT18 in phase with *Pura* transcription (Fig. 1B).

**Figure 5.**
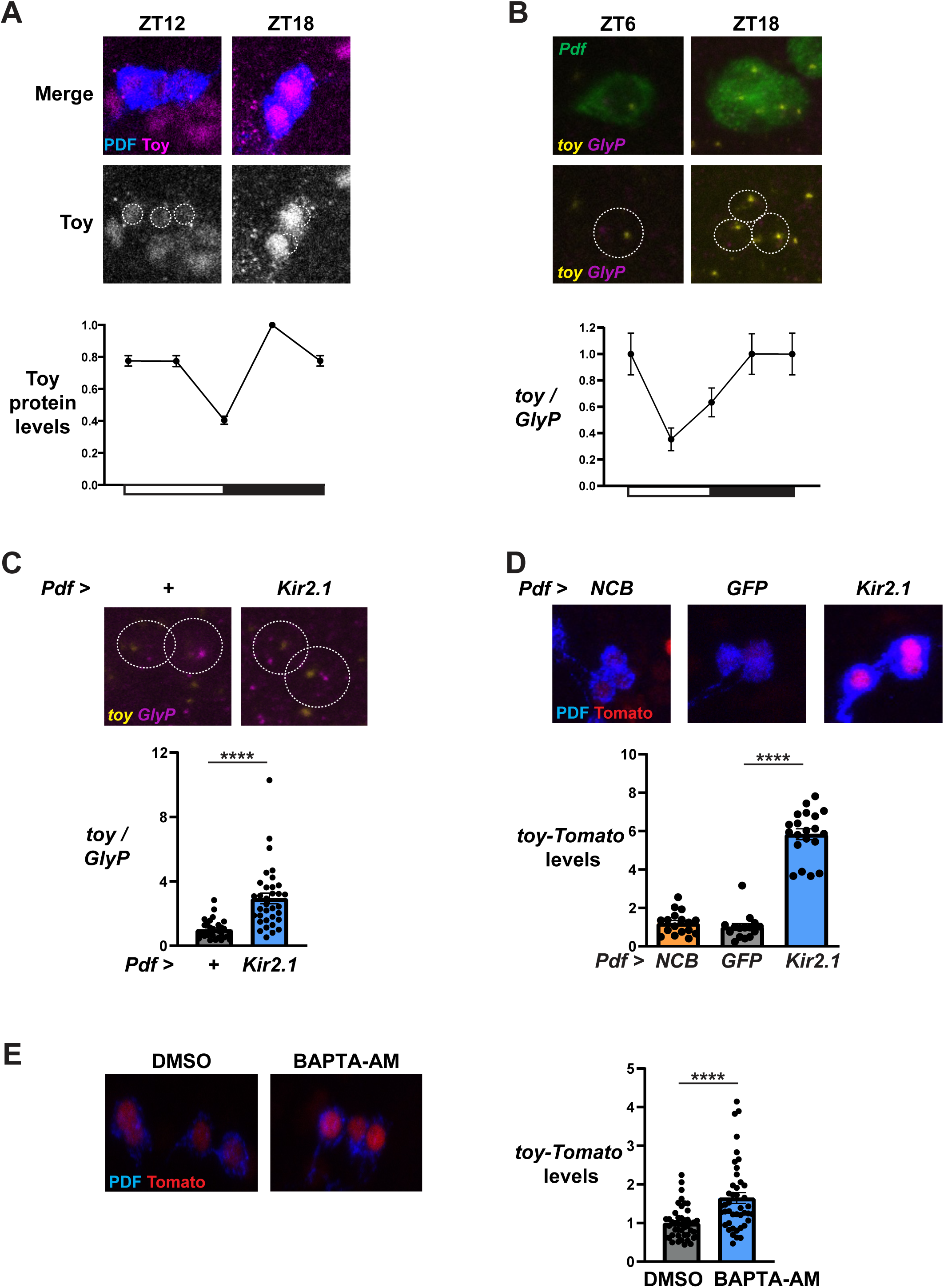
*toy* is regulated in LN_v_s by neuronal inactivity. **(A) Rhythms in Toy protein levels in LN_v_s.** Control (*y w*) larvae were entrained in LD cycles at 25°C, dissected at the times shown and stained with antibodies to Toy (purple) and PDF to mark LN_v_s (blue). Top: Representative confocal images from ZT12 and ZT18. Middle: Single channel images show Toy signals in grayscale with dashed lines circling the areas used for quantification. Bottom: Toy signals in LN_v_s were quantified, normalized to the average of the peak time point and plotted. Data from ZT0 are replotted at ZT24. Error bars show SEM. Toy protein levels vary with time (one-way ANOVA, p < 0.0001). The graph is an average of 2 independent experiments with 15-20 LN_v_s from 5 brains in each experiment. **(B) *toy* transcription is rhythmic in LN_v_s.** Control (*y w*) larvae were entrained in LD cycles at 25°C, dissected at the times shown and *in situ* hybridization performed using intron probes for *toy* (yellow) and *GlyP* (magenta), and exon probes for *Pdf* (green). Top: Representative confocal images from ZT6 and ZT18 showing all 3 channels. Middle: The same images as above, but with the green channel omitted to visualize intronic signals in the nucleus more clearly. Bottom: Time course showing the intensity of *toy* intron in situ hybridization signal divided by *GlyP* intron signal for all LNvs in a hemisphere and normalized to ZT24. The data from ZT24 is replotted at ZT0. Error bars show SEM. *toy* / *GlyP* levels vary with time (one-way ANOVA, p < 0.05). One-way ANOVA followed by Tukey’s multiple comparisons test to compare timepoints show that *toy* / *GlyP* intensity at ZT6 was less than that at ZT0 and ZT18 (p < 0.05). Data are from two independent experiments with at least 3 brains per timepoint in each experiment, and 3-4 LN_v_s per hemisphere. **(C) *toy* transcription is activated by hyperpolarization.** Top: Representative confocal images of *in situ* hybridizations using intronic probes for *toy* (yellow) and *GlyP* (magenta), and exonic probes for *Pdf* (outlined in white circle) from either *Pdf-Gal4* / + control larvae or *Pdf-Gal4* expressing *UAS-Kir2.1* (*Pdf > Kir2.1*) larvae at ZT12. Bottom: *toy / GlyP* ratios, with each dot representing a single LNv, and normalized to the average of control LN_v_s. Error bars show SEM. Unpaired t test shows *toy* intron intensity is higher when LN_v_s express Kir (p < 0.0001). Data is from one experiment with at least 6 brains per genotype. We saw a very similar result in an independent experiment without including *GlyP* as a loading control. **(D) A *toy* transcriptional reporter gene is activated in hyperpolarized LN_v_s.** *toy-Tomato* levels were measured in LN_v_s from larvae with *Pdf-Gal4* expressing *UAS-NaChBac* (*Pdf > NCB*), *UAS-GFP* (*Pdf > GFP*) or *UAS-Kir2.1* (*Pdf > Kir2.1*). Larval brains were dissected at ZT12, stained, quantified and the data normalized as in Fig. 1C. Top: representative confocal images with PDF (blue) to mark LN_v_s and Tomato (red). Bottom: *toy-Tomato* levels are higher in *Pdf > Kir2.1*-expressing LN_v_s (blue bar) than in control *Pdf > GFP* LN_v_s (grey bar). Student’s *t*-test, p < 0.0001. There is no difference in *toy-Tomato* levels between *Pdf > NCB* (orange bar) and control LN_v_s (Student’s *t*-test, p > 0.05). Error bars show SEM. The graph is an average of 2 independent experiments with 15-20 LN_v_s from 5 brains in each experiment. **(E) *toy* transcription is increased by chelating intracellular Ca^2+^.** Left: representative confocal images of larval brains showing PDF (blue) and Tomato (red)*. toy*-tomato larval brains were dissected at ZT8 and incubated in Schneider’s media containing either 0.1% DMSO or 20μM BAPTA-AM for 4 hours before fixing and staining. Right: Each dot represents *toy-*Tomato intensity in a single LNv. Data were normalized to levels in the control vehicle (DMSO). Bars represent average *toy-*Tomato intensity. Errors bars show SEM. Unpaired t test shows *toy-*Tomato levels are higher when incubated in BAPTA-AM (p < 0.0001). Data show the average of two independent experiments with at least 4 brains per experiment.

Next we tested if *toy* transcription is regulated in LN_v_s using *in situ* hybridization with probes that detect *toy* intronic sequences. These probes label 1-2 spots in the nucleus that represent nascent transcripts and thus reflect active transcription.^52^ We also included probes that recognize the first intron of *Glycogen phosphorylase* (*GlyP*), which is highly transcribed in LN_v_s and many other neurons (Fig. 5B), and the *Pdf* exon to label LN_v_s. We then calculated the ratio of intensity of the *toy* and *GlyP* spots to normalize *in situ* hybridization signals between brains.

The data in Fig. 5B show that *toy* transcription is rhythmic in larval LN_v_s, with levels decreasing between ZT0 and ZT6, and then increasing from ZT6 to ZT18. To test if *toy* transcription is regulated by neuronal inactivity, we measured *toy* transcription using the intronic *in situ* hybridization probes in control LN_v_s and in LN_v_s constitutively expressing the *UAS-Kir2.1* transgene. We found that expressing Kir2.1 in LN_v_s induces *toy* transcription (Fig. 5C).

We also decided to make a *toy* transcriptional reporter gene. Two overlapping *toy* enhancers (R9G08 and R9G09) appear to drive expression in LN_v_s^23^ and the 1.1kb overlapping region is highly conserved across *Drosophila* species. We generated a *toy* transcriptional reporter by fusing the *toy* enhancer R9G08 plus 87bp from R9G09 to an *Hsp70* basal promoter and the destabilized tdTomato-NLS reporter gene. We then measured *toy-Tomato* levels at ZT12 in LN_v_s expressing Kir2.1. The data in Fig. 5D confirm that *toy* transcription increases in *Kir2.1*-expressing LN_v_s compared to control LN_v_s. Therefore transcription of the *Pura* regulator Toy is activated in response to decreases in neuronal activity in LN_v_s.

### A mechanism to activate *toy* transcription

The data presented so far show that decreasing neuronal activity increases transcription of *toy*, and that Toy activates *Pura* expression – an effector gene for neuronal plasticity.^13^ This “inactivity-regulated gene pathway” parallels the activity-regulated program, in which increasing neuronal activity activates transcription of activity-regulated genes, some of which encode transcription factors that then transcribe effector genes for plasticity.^53^ Indeed, increasing s-LNv firing leads to rapid structural plasticity and expansion of s-LNv projections^14^, as well as transcription of activity-regulated genes like Hr38^16^ (Fig. 2A), indicating that this pathway is active and functional in LNvs. Thus it seems important to ensure that either the activity-regulated gene <u>o</u>r the inactivity-regulated gene programs is active in a cell at one time. How could this be achieved?

Neuronal firing increases intracellular Ca^2+^ and activates enzymes like CaMKII to phosphorylate transcription factors such as CREB which then increase transcription of activity-regulated genes.^53^ We wondered if *toy* – as an inactivity-regulated gene – would be activated by reduced intracellular Ca^2+^ levels that could come from periods of decreased neuronal activity.

Intracellular Ca^2+^ levels in adult s-LNvs follow a 24 hour rhythm in constant darkness (DD) with lowest intracellular Ca^2+^ levels from shortly before subjective dusk (∼CT9) onwards.^54^ This is consistent with the timing of when *toy* transcription starts to increase in larval LNvs (Fig. 5B).

To test if reducing intracellular Ca^2+^ levels activate *toy* transcription, we dissected *toy-Tomato* larval brains at ZT8 and incubated them for 4 hours in media to which we added either vehicle (DMSO) or the cell-permeable Ca^2+^ chelator, BAPTA-AM in DMSO. Brains were then fixed and stained. The data in Figure 5E shows that explanted brains incubated in BAPTA-AM show a modest but reproducible increase in levels of the *toy-Tomato* transcriptional reporter gene.

Therefore reducing intracellular Ca^2+^ levels is sufficient to increase *toy* transcription, which is the opposite of how canonical activity-regulated genes respond to intracellular Ca^2+^.

### Toy regulates s-LNv structural plasticity

Finally, we tested if *toy* is important in s-LNv structural plasticity. We first induced *toy* overexpression using a *UAS-toy* transgene controlled by a combination of the *Pdf-Gal4* driver and the *tubulin-Gal80^ts^*transgene to block Gal4 activity at low temperatures. Flies also contained a *Pdf-RFP* transgene^28^ to visualize s-LNv projections. Flies were raised at 18°C and then entrained in LD cycles at 18°C, before shifting to 29°C for 4 hours. Brains were dissected at CT2, which is 2 hours after subjective lights on in DD. Flies were kept in DD to prevent any light-induced neuronal firing at dawn. s-LNv projections were then imaged and quantified using a custom MATLAB script.^13^ Fig. 6A shows that control s-LNvs, which went through the same temperature and light cycles as the experimental s-LNvs, have normal expanded s-LNv projections at CT2. In contrast, inducing *toy* expression for just 4 hours blocked s-LNv structural plasticity at dawn, with projections in a dusk-like retracted state.

**Figure 6.**
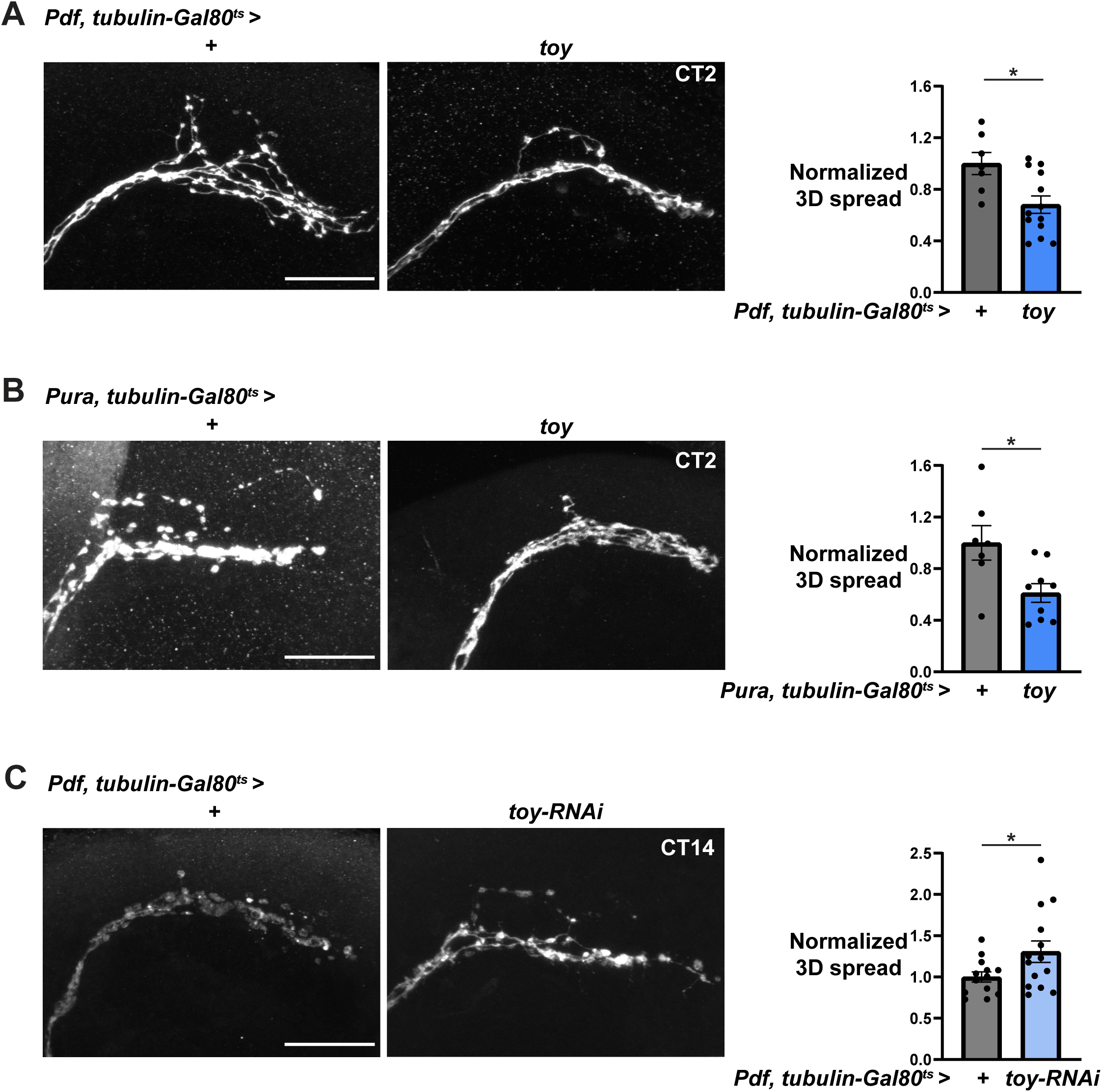
Toy regulates the structural plasticity of adult s-LN_v_s. **(A) Transiently over-expressing *toy* with *Pdf-Gal4* at dawn blocks the expansion of s-LNv projections.** Left: Representative confocal images of s-LN_v_ projections from flies containing *Pdf-Gal4* and a temperature-sensitive Gal80 transgene (*tubulin-Gal80^ts^*) crossed to either *y w* control flies or flies with a *UAS-toy* transgene. *toy* overexpression was induced by shifting flies from 18°C to 29°C for 4 hours at ZT22 and dissecting at CT2. Flies also contained a *Pdf-RFP* transgene, and s-LN_v_ dorsal projections were visualized by an antibody to RFP. Scale bar = 25 µm. Right: Quantification of the 3D spread of s-LN_v_ projections. Bars represent the average 3D spread normalized to control s-LN_v_s. Each dot represents the 3D spread of one set of s-LN_v_ projections from one hemisphere. Only one hemisphere was quantified per brain. Error bars show SEM. s-LN_v_ projections in flies over-expressing *toy* in all LN_v_s have reduced 3D spread compared to control flies (unpaired t-test, p = 0.01). The data are from at least 7 brain hemispheres from 2 independent experiments. **(B) Transiently over-expressing *toy* with an s-LN_v_ specific driver blocks the expansion of s-LN_v_ projections at dawn.** Left: Representative confocal images of s-LN_v_ projections as in A except that *Pura-Gal4* was used for *toy* overexpression, which was induced by shifting flies from 18°C to 29°C at ZT12 and dissecting 14 hours later at CT2. s-LN_v_ projections were visualized via a *Pdf-RFP* transgene as in A. Scale bar = 25 µm. Right: Quantification of the 3D spread of s-LN_v_ projections as in A. s-LN_v_ projections in flies over-expressing *toy* in s-LN_v_s have reduced 3D spread compared to control flies (unpaired t-test, p < 0.05). The data are from at least 7 brain hemispheres from 2 independent experiments. **(C) Transiently reducing *toy* levels expands s-LN_v_ projections at dusk.** Left: Representative confocal images as in A of s-LN_v_ projections from flies containing *Pdf-Gal4* and *tubulin-Gal80^ts^* crossed to either *y w* control flies or flies with a *UAS-toy-RNAi* transgene. *toy-RNAi* expression was induced by shifting flies from 18°C to 29°C at CT0 for 14 hours and dissecting at CT14. Scale bar = 25 μm. Right: Quantification of the 3D projections as in A. s-LN_v_ projections in flies expressing *toy-RNAi* in LN_v_s have increased 3D spread compared to control flies (unpaired t-test, p < 0.05). The data are from at least 12 hemispheres from 2 independent experiments.

*Pdf-Gal4* is expressed in both the small- and large-LNvs.^55^ To confirm that the effect of *toy* on s-LNv plasticity is cell-autonomous, we repeated the *toy* overexpression experiment, but this time using a *Pura-Gal4* driver, which expresses in s-LNvs but not in l-LNvs.^56^ The data in Fig. 6B show that overexpressing *toy* using *Pura-Gal4* and *tubulin-Gal80^ts^*for 14 hours blocked the normal expansion of s-LNv projections at dawn (CT2).

We also tested if *toy* is required for s-LNv projections to retract at subjective dusk (CT14). We used a *UAS-toy-RNAi* transgene and induced expression for 14 hours starting at CT0 using the *Pdf-Gal4* and *tubulin-Gal80^ts^* combination. The data in Fig. 6C show that expressing *toy-RNAi* in LNvs blocks the full retraction of s-LNv projections at dusk. The variability in the dataset probably comes from incomplete knockdown of *toy* that differs between flies and/or additional post-translational regulation of s-LNv retraction via the FMRP RNA-binding protein.^57^

## Discussion

Transcription of genes such as *c-fos* is rapidly induced in mammalian neurons in response to increased neuronal activity. Many of these immediate early genes encode transcription factors that regulate a second set of genes and overall these two waves of gene expression stabilize changes in the excitability and connectivity of neurons in response to firing.^58^ Here we describe a complementary gene expression program that is activated in response to neuronal inactivity and regulates neuronal plasticity. Using *Drosophila* LN_v_s, we showed that transcription of the gene encoding the Pax-6 transcription factor Toy is activated by reduced neuronal activity and decreased intracellular calcium. Toy then activates transcription of *Pura*, a plasticity gene which encodes a Rho1 GEF that increases Rho1 activity to retract s-LN_v_ projections and reduce their signaling potency.^13^

There are many genes whose transcription increases in response to firing. So far, we only know of 2 genes – *toy* and *Pura* – whose transcription is induced by neuronal inactivity in LNvs. While it may be premature to call this a gene expression program, induction of the transcription factor Toy as a putative first responder makes it likely there are multiple target genes. Furthermore, we identified many genes whose expression was activated by long-term expression of Kir2.1 in LN_v_s.^15^ However, we do not yet know if these genes are Toy targets or that their RNA levels are regulated transcriptionally.

### Are inactivity-regulated genes a general property of neurons?

Singing induces the expression of hundreds of genes in the brain of zebra finches, and simultaneously represses hundreds of other genes.^59^ Some of these latter genes could be inactivity-regulated genes. In addition, blocking neuronal activity in cultured mouse cortical neurons induced the expression of many genes.^60^ Furthermore, neuronal hyperpolarization has an instructive role in mouse neocortex development because precociously hyperpolarizing neuronal progenitors changed their transcriptional profile and advanced their developmental trajectory.^61^ Together, these data suggest that a gene expression program that responds to neuronal inactivity exists beyond LN_v_s. Mammalian neurons are certainly able to recognize reductions in neuronal inactivity because phosphorylation of Pyruvate dehydrogenase increases as neuronal activity decreases.^62^ It will be very interesting to know if inactivity-driven gene expression is a general property of neurons.

### A mechanism for expression of inactivity-regulated genes

If one function of this inactivity-regulated gene expression program is to retract projections, then it makes sense that activity-regulated and inactivity-regulated gene expression programs are not active at the same time. Transcription factors from each program regulate *Pura* expression via a small region of the *Pura* enhancer, perhaps even competing for binding to overlapping sites on DNA (see Fig. 7). The intracellular signaling pathways activated by neuronal firing that rapidly lead to induction of gene expression are well-understood.^40,63^ However, we do not yet fully understand how *toy* transcription is activated in response to neuronal inactivity. Activity-regulated and inactivity-regulated genes responding to opposite changes in intracellular Ca^2+^ levels fits with the idea that LN_v_s should not try to expand and retract their projections at the same time. Thus one signaling molecule triggering both gene expression programs – but at opposite levels – would coordinate the two programs since it is impossible for intracellular Ca^2+^ concentrations to simultaneously be high and low in one cell. However, we speculate that the kinetics of these two gene expression programs are different. While it seems advantageous for a neuron to rapidly respond to firing by strengthening synapses, a neuron should perhaps only start to induce a synaptic retraction or pruning program with much slower kinetics. In this way, the threshold for “low” intracellular Ca^2+^ to activate inactivity-regulated genes may only be reached after a longer time – perhaps only after prolonged inactivity. Testing this will be helped by developing tools with bright enough fluorescence to follow transcription in vivo in real time.

**Figure 7.**
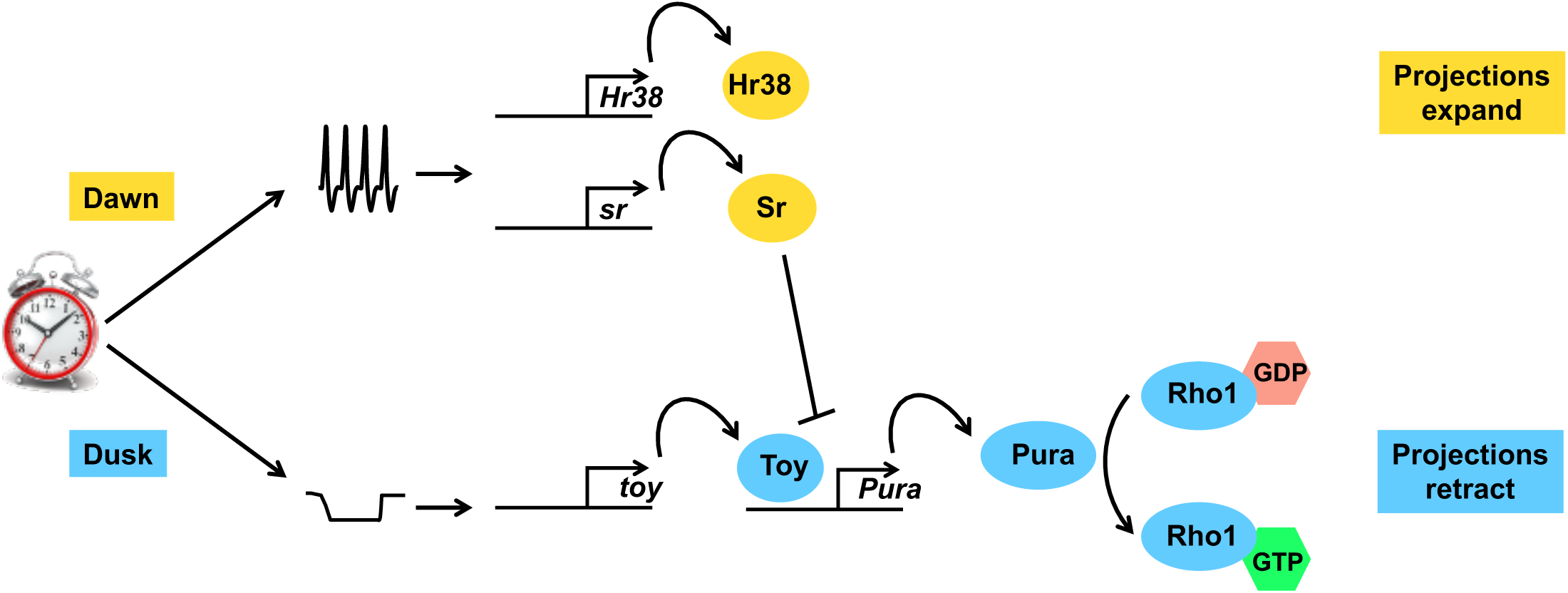
Activity-regulated and inactivity-regulated gene expression programs in s-LN_v_s. The circadian clock in s-LN_v_s leads to increased neuronal firing at dawn which activates the canonical activity-regulated gene pathway (yellow) and promotes structural plasticity by expanding s-LN_v_ projections. At least one activity-regulated gene (Sr) represses *Pura* transcription. Reduced neuronal activity and/or hyperpolarization of LN_v_s around dusk induces *toy* transcription. Toy then activates transcription of *Pura*. The Rho1 GEF Pura then increases Rho1 activity to promote plasticity by retracting s-LN_v_ projections (blue pathway).

Rhythmic and predictable plasticity is not unique to s-LNvs. There are 4-5 day rhythms in dendritic spine number and synaptic density in the hippocampus of female mice which follow the oestrous cycle.^64–66^ The molecular mechanisms may even be similar: the Rho Guanine nucleotide Exchange Factor ARHGEF5 is one of the top upregulated genes during dioestrus when dendritic spines are retracted.^66^ ARHGEF5 increases RhoA activity^67^ and thus could act in the same way that Pura increases Rho1 activity to retract s-LNv projections.^13^ It will be interesting to identify the molecular controls on ARHGEF5 expression. Another example of rhythmic plasticity comes from the synaptic downscaling that seems to reduce the overall connectivity of the brain during sleep ^7,8^. We speculate that a similar gene expression program to the one described here could help restore parts of the brain to a similar level of connectivity as on waking the previous day.

## Figure Legends

**Figure S1 (related to Fig. 1).**
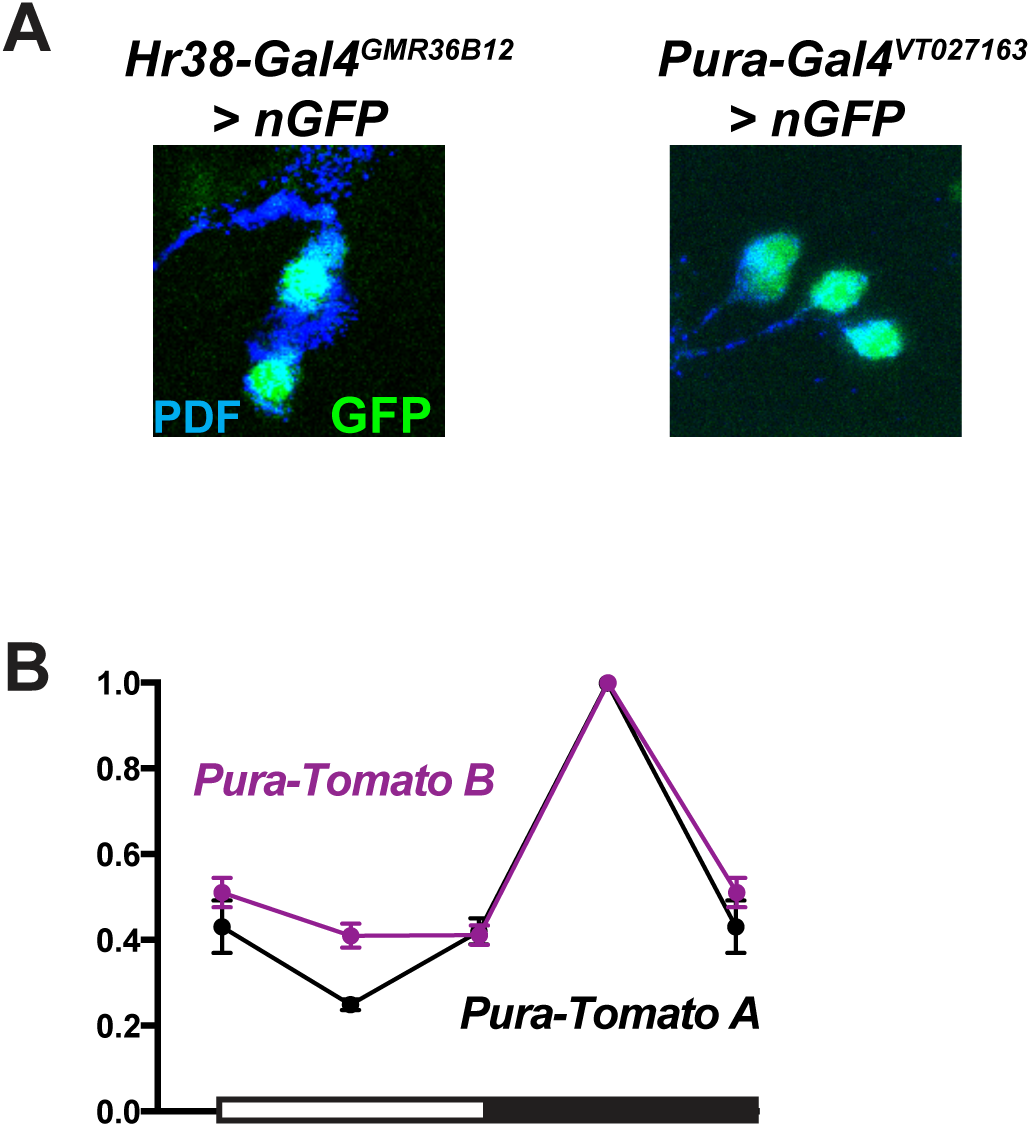
*Hr38* and *Pura* expression in LN_v_s. **(A) Identifying *Hr38* and *Pura* enhancers that express in LN_v_s.** Larvae with *UAS-nGFP* and either *Hr38-Gal4^GMR36B^*^12^ or *Pura-Gal4^VT^*^027163^ transgenes were entrained in LD cycles at 25°C. Third instar larval brains were dissected and stained with antibodies to PDF to mark LN_v_s (blue) and GFP (green). Left: Representative confocal image of *Hr38-Gal4^GMR36B^*^12^ expressing GFP at ZT2. Right: Representative confocal image of *Pura-Gal4^VT^*^027163^ expressing GFP at ZT15. **(B) *Pura-Tomato A* and *Pura-Tomato B* have similar phase rhythms in larval LN_v_s.** Transgenic larvae with either *Pura-Tomato A* (black line) or *Pura-Tomato B* (purple) were entrained in LD cycles at 25°C and brains dissected at the times shown. Staining, quantification and normalization are as in Fig. 1B with data from ZT0 replotted at ZT24. *Pura-Tomato A* and *B* levels vary with time in LN_v_s (one-way ANOVA, p < 0.0001). Error bars show SEM. The graph is an average of 2 independent experiments with 15-20 LN_v_s from 5 brains in each experiment.

**Figure S2 (related to Fig. 2).**
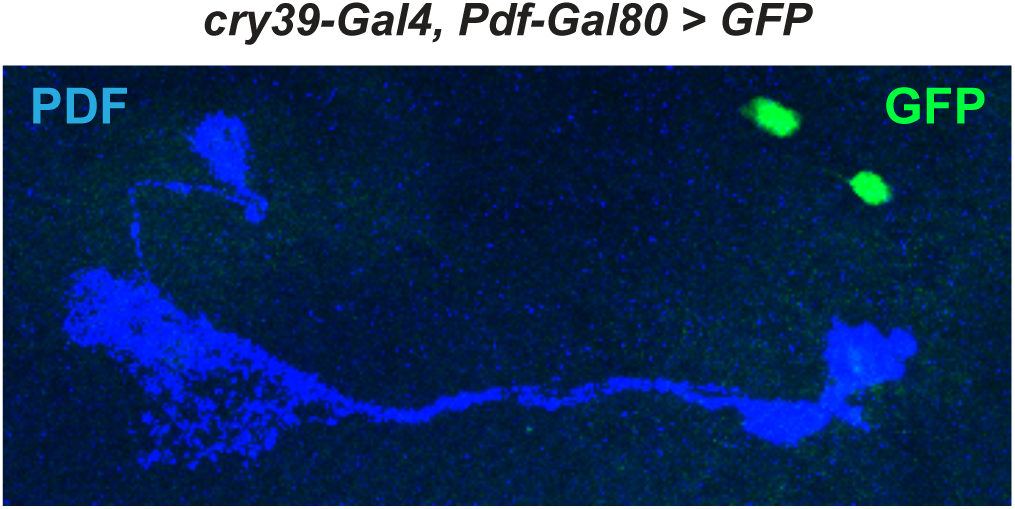
*Pdf-Gal80* blocks *cry39-Gal4* activity in LNvs. Larvae with *cry39-Gal4*, *Pdf-Gal80* expressing *UAS-GFP* transgenes in DN1s were raised at 22°C in LD cycles. Larvae were shifted to 30°C at ZT8 for 4hr before dissection at ZT12 – see timeline in Fig. 2C. Larval brains were stained with antibodies to GFP to mark DN1s (green), PDF to mark LN_v_s (blue). Representative confocal image shows the temperature-independent *Pdf-Gal80* transgene ^37^ blocks Gal4 activity in LN_v_s and prevents GFP expression in LN_v_s.

**Figure S3 (related to Fig. 3).**
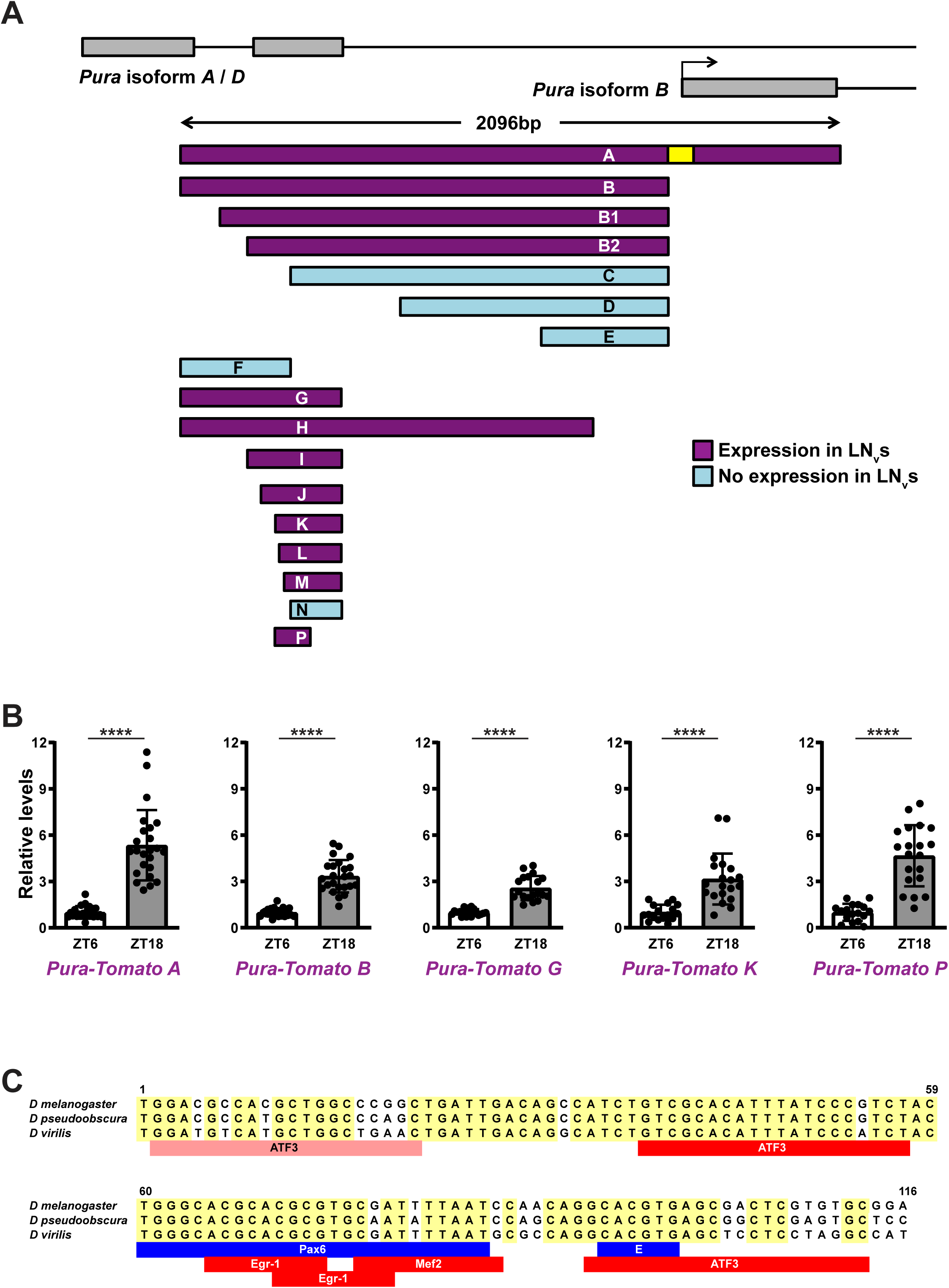
Dissecting the *Pura* enhancer identified a minimal fragment that expresses rhythmically in LN_v_s. **(A) A minimal *Pura* enhancer for LN_v_ expression.** Map of the *Pura* regulatory region showing exons (grey boxes), introns (black lines) and the transcription state site (TSS) with a black arrow. *Pura* enhancers (purple or blue) were fused with the *Pura* basal promoter (−50bp to +50bp of TSS, yellow) to drive Tomato expression. This map shows the *Pura* enhancers we tested including *Pura* enhancer A, B and P which were shown in Fig. 3A. Larvae with a single *Pura-Tomato* transgene were entrained in LD cycles at 25°C. Larval brains were dissected at the peak expression time ZT18, stained and scanned as in Figure 1B. *Pura-Tomato* transgenes expressed in LN_v_s (A, B, B1, B2, G, H, I, J, K, L, M and P) are shown in purple and reporter genes with undetectable levels in LN_v_s (C, D, E F and N) are in blue. **(B) *Pura* enhancers drive rhythmic expression in LN_v_s.** Larvae with a *Pura-Tomato* transgene (A, B, G, K or P) were entrained in LD cycles at 25°C. Larval brains were dissected at ZT6 and ZT18, stained and quantified as in Fig. 1B. *Pura-Tomato* levels were normalized to the average level at ZT6 in each experiment. Levels of all of these *Pura-Tomato* transgenes were lower at ZT6 than ZT18 (Student’s *t*-test, p < 0.0001). Error bars show SEM. **(C) Transcription factor binding sites in *Pura enhancer P* are conserved across *Drosophila* species.** The 116bp DNA sequence of *Pura enhancer P* from *D. melanogaster* was compared with sequences from *D. pseudoobscura* and *D. virilis*. Identical bases are shaded in yellow. Locations of potential transcription factor binding sites were marked by red (ATF3, Egr-1 and Mef2) or blue boxes (Pax6 and E-box) underneath the DNA sequences. For the 3 potential ATF3 binding sites found in *D. melanogaster*, the one that was not conserved in either *D. pseudoobscura* or *D. virilis* was marked as a pink box; the two conserved in *D. pseudoobscura* but not in *D. virilis* were marked as a red box.

**Figure S4 (related to Fig. 4).**
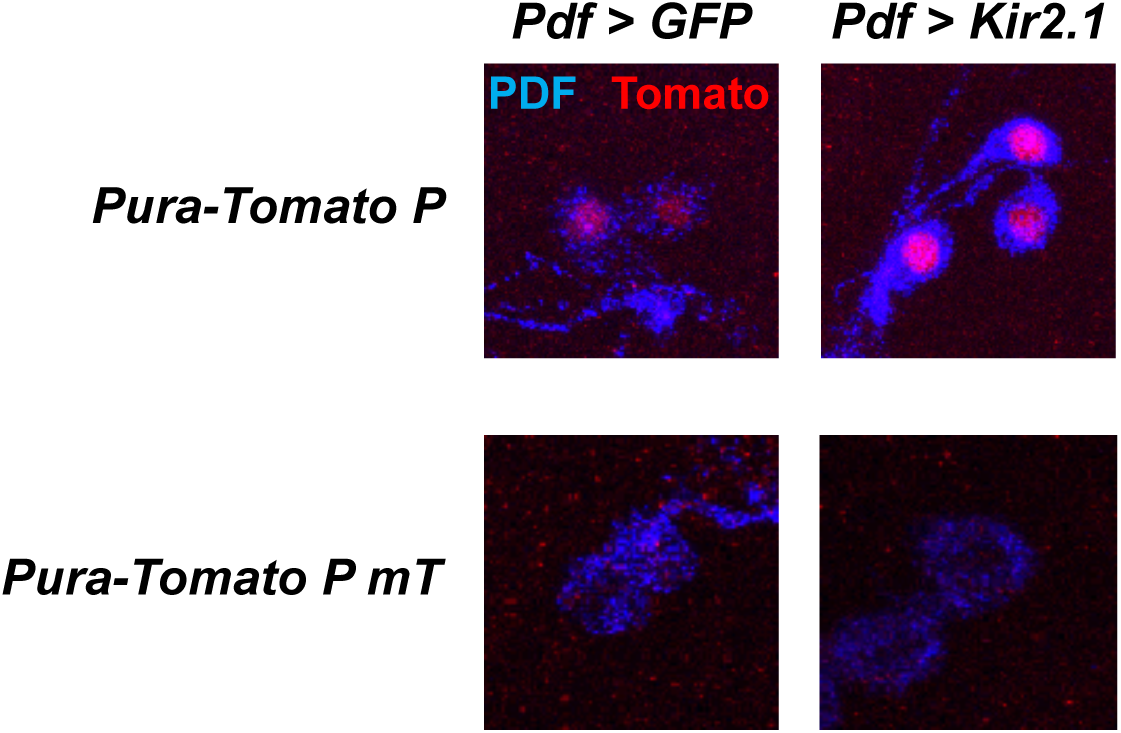
*Pura-Tomato P mT* is undetectable in Kir2.1-expressing LN_v_s. Larvae with *Pdf-Gal4* and either *Pura-Tomato P* or *Pura-Tomato P mT* transgenes were entrained in LD cycles at 25°C. Control larvae expressed a *UAS-GFP* transgene in LN_v_s (*Pdf > GFP*, left) and experimental larvae expressed a *UAS-Kir2.1* transgene in LN_v_s (*Pdf > Kir2.1*, right). Larval brains were dissected, stained and scanned as in Fig. 1B. Representative confocal images are shown with PDF to mark LN_v_s (blue) and Tomato (red). Top: *Pura-Tomato P* in LN_v_s at ZT12. Bottom: *Pura-Tomato P mT* in LN_v_s at ZT18, the peak of *Pura* expression in LN_v_s.

**Table S1 (related to Fig. 4)**

**Details of the RNAi screen to identify the transcription activator of *Pura*.**

Larvae with *Pura-Tomato B*, *Pdf-Gal4*, *UAS-Dcr-2* and one of 110 *UAS-shRNA* transgenes targeting transcription factors or a control transgene *UAS-GFP* (*GFP*) were entrained in LD cycles at 25°C. Larval brains were dissected at ZT18, stained and quantified as in Fig. 1B. Tomato signals in single LN_v_s were normalized to the mean of the GFP-expressing control LN_v_s. Different columns show the mean, quantified cell number (N) and SEM of *Pura-Tomato B* levels in LN_v_s with each *UAS-shRNA* transgene.

This table is available on request.

## Methods

### 1. Cloning strategies

We modified pBPGUw ^69^, Addgene) to create a vector to make fluorescent reporter genes for different enhancers. pBPGUw contains a Gateway cassette to easily insert enhancers through the LR reaction upstream of the synthetic basal promoter DSCP. pBPGUw also contains a *PhiC31 attB* sequence for site-specific genome insertion and a mini *white^+^* gene to select transformants. We PCR-amplified DNA for codon-optimized *tdTomato*, 3 copies of nuclear localization sequence, a *PEST* sequence and a *SV40* 3’UTR ^25^. PCR products were TOPO cloned to the pCRXL-TOPO vector (Life Technologies), digested out with KpnI and SpeI. This insert was ligated to KpnI / SpeI-digested pBPGUw which removes the *Gal4* sequence. We called the resulting plasmid pZT.

Since Pfeiffer et al. ^26^ and our preliminary data showed that the *DSCP* basal promoter is relatively weak, we replaced *DSCP* with the *Hsp70* basal promoter. We also added an intervening sequence (IVS) and a synthetic 5’UTR (*Syn21*) to increase translational efficiency as in ^26,27^. Since there were no restriction sites available for direct cloning, a DNA fragment containing the Hsp70 basal promoter, IVS, *Syn21* 5’UTR and part of tdTomato cDNA was synthesized by Integrated DNA Technologies using the sequences from ^26,27^. This DNA fragment was flanked by FseI and SbfI sites. The fragment was amplified by PCR, TOPO cloned to pCR8GWTOPO, and digested by FseI and SbfI to remove it from the vector. This fragment was inserted into pZT digested with FseI and SbfI to create pZT2.0. The DNA sequence was verified by Sanger sequencing (Genewiz).

For *Hr38-Tomato*, we used PCR to amplify the 4kb of *Hr38* regulatory DNA in *Hr38^GMR36B^*^12^*-Gal4* ^23^ from fly genomic DNA. This DNA was TOPO cloned to pCR8GWTOPO vector to create pCR8GWTOPO-*Hr38 enhancer*. Then we performed an LR reaction (Gateway^TM^ LR Clonase^TM^ II Enzyme mix, Thermo Scientific) to insert the *Hr38* enhancer into pZT2.0 to make *Hr38-Tomato*.

All *Pura-Tomato* transgenes were driven by an endogenous *Pura* basal promoter (−50bp to +50bp of the transcription start site of *Pura* isoform B). We modified pZT2.0 to either remove the HSP70 basal promoter (pZT2.3) or replace it with the *Pura* basal promoter (pZT2.4). To make pZT2.3, we digested pZT2.0 with FseI and BglII and ligated a 22bp synthetic fragment with FseI and BglII sticky ends. Then we used PCR to amplify the *Pura* enhancers A, B, B1, B2, C, D and E from fly genomic DNA. These fragments contained the *Pura* basal promoter. They were cloned into pCR8GWTOPO and LR reactions were used to insert these enhancers into pZT2.3 to generate *Pura-Tomato* transgenes.

pZT2.4 contains the *Pura* basal promoter in the backbone. We synthesized an 114bp fragment containing the *Pura* basal promoter with an FseI site and a BglII site on the 5’end and 3’ end respectively. We cloned the 114bp basal promoter fragment into pCR8GWTOPO and cut with FseI and BglII to make the insertion. We also cut pZT2.0 with FseI and BglII to create the backbone and ligated it with the *Pura* basal promoter insertion. This generated plasmid pZT2.4. We used PCR to amplify *Pura* enhancer fragments F, G, H, I, J, K, L, M, N and P from fly genomic DNA, which do not include the *Pura* basal promoter. Then we used TOPO cloning to insert each one to pCR8GWTOPO and used LR reactions to insert these enhancers into pZT2.4 to generate *Pura-Tomato* transgenes.

To make *Pura-Tomato P mE*, we changed the E-box sequence CACGTG in *Pura enhancer P* to CA**GC**TG (mutations in bold) according to ^42^ and synthesized the mutated 116bp DNA sequence (Integrated DNA Technologies). We cloned the synthesized DNA fragment into the pCR8GWTOPO vector to create pCR8GWTOPO-*Pura enhancer P mE* and then performed an LR reaction to insert *Pura enhancer P mE* to pZT2.4 to generate *Pura-Tomato P mE*.

To make Pura-Tomato PmT, we mutated the sequence TCTACTGGGCACGCACGC in *Pura enhancer P* that contains the Toy binding site to T**T**T**C**C**G**G**A**G**T**A**T**G**T**A**T**G**T** (mutations in bold) and synthesized the mutated 116bp DNA sequence (Integrated DNA Technologies). We cloned the synthesized DNA fragment into the pCR8GWTOPO vector to create pCR8GWTOPO-*Pura enhancer P mT* and then performed an LR reaction to insert *Pura enhancer P mT* to pZT2.4 to generate *Pura-Tomato P mT*.

For *toy-Tomato*, we used PCR to amplify a 3.6kb *toy enhancer* DNA fragment from fly genomic DNA and cloned it into the pCR8GWTOPO vector to create pCR8GWTOPO-*toy enhancer*. Then we performed an LR reaction to insert the *toy enhancer* to pZT2.0 to make *toy-Tomato*.

### 2. Generating transgenic animals

All plasmids containing reporter transgenes were prepared using Qiagen Plasmid Mini Kit and were sent to Genetivision Corporation for injection. All reporter transgenes were inserted to a specific site using PhiC31 recombinase mediated methods ^70^. All transgenes were inserted to VK37: (2L)22A3 ^71^. Since all of the transgenes contain the mini *w^+^* selection marker, plasmids were injected into *w* mutant embryos to make transgenic lines and crossed to *w* mutant flies to screen for *w^+^*transgenic flies. Red-eyed flies were then balanced to make stock lines.

### 3. Immunofluorescence staining

Larvae and flies developed at 25°C in LD cycles unless specifically noted. Brains of *Drosophila* 3^rd^ instar wandering larvae were dissected at specific times of day in phosphate-buffered saline (PBS) and immediately fixed in fresh 4% formaldehyde in PBS for 20 min at room temperature. Brains were then permeabilized by washing twice in PBS with 1% Triton X-100 for 10 min and then blocked in PBS with 0.5% Triton X-100 and 10% normal horse serum (VECTOR S-2000) for at least 30 min at room temperature. Brains were incubated in PBS with 0.5% Triton X-100 and 10% normal horse serum and primary antibodies overnight at 4°C. Brains were rinsed once and then washed 10 min for 3 times in PBS with 0.5% Triton X-100 before incubating with secondary antibodies in PBS with 0.5% Triton X-100 and 10% normal horse serum either at room temperature for 1 hr or overnight at 4**°**C. Brains were rinsed once and then washed 10 min for 3 times in PBS with 0.5% Triton X-100. Brains were soaked in 50% glycerol in PBS or SlowFade Gold antifade reagent (Invitrogen S36936) for 20 min before mounting.

Antibody list:

Primary antibodies:

PDF: “PDF C7”, Developmental Studies Hybridoma Bank, mouse, used at 1:100 RFP: “PM005”, MBL, rabbit, used at 1:500

Toy: gift from Claude Desplan, previously unpublished, rabbit, used at 1:500 GFP: “NB100-62622”, Novus Biologicals, sheep, used at 1:500

Secondary antibodies:

Alexa Fluro 488 donkey anti mouse (Life Technologies, A21202) Alexa Fluro 488 donkey anti sheep (Life Technologies, A11015) Alexa Fluro 555 donkey anti rabbit (Life Technologies, A31572) Alexa Fluro 647 donkey anti rabbit (Life Technologies, A31573) Alexa Fluro 647 donkey anti mouse (Life Technologies, A31571) All used at 1:500.

### 4. Confocal Microscopy and Image Analysis

Brains were scanned as single stack images with 1μM in depth (1024 X 1024 or 1024 X 256 resolution) under the 20X objective lens of Leica SP5 confocal microscope with 2X or 4X digital zoom in. For quantification, a stack of images (usually 10-13 images) was projected together on the Z-axis. Mean fluorescence intensity in the soma of single neurons were quantified using FIJI with background subtracted. Background levels were determined by the average of mean fluorescence intensity of 10 regions with the same size randomly selected in the same brain lobe.

### 5. Bioinformatics analysis to search for transcription factor binding sites

We first used JASPAR CORE Vertebrata [jaspar.genereg.net, ^72^] with default settings to predict CREB and Mef2 binding sites in the *Hr38* and *Pura* enhancers. We chose the database of vertebrate transcription factors binding sites because it includes more transcription factors (1045 for vertebrates vs 266 for insects). Then we used MATCH analysis from TRANSFAC ^39^ with default settings to identify potential CREB, Mef2, Egr1, Atf3 and Pax6 binding sites as well as E-boxes. We used the vertebrate transcription factor PWMs for the same reason as above. In addition, the vertebrate transcription factor PWM database is better annotated for Pax6 as it includes the Paired domain binding sequence as well as the Homeodomain binding sequence, while the insect PMW only has the Homeodomain binding sequence.

### 6. Conservation analysis

We used the bioinformatics tool phastCons analysis in the UCSC genome browser ^38^ to identify DNA sequences in the *Pura enhancer P* that are conserved in *D. pseudoobscura* and *D. virilis*. Then we used MATCH analysis to predict transcription factor binding sites in these regions in *D. pseudoobscura* and *D. virilis* as above.

### 7. Statistics

All data points were normalized to the mean of the control group unless specified in figure legends. Error bars in all figures show mean ± SEM. Sample size (n) of all experiments is indicated in figure legends. All experiments with LN_v_s measured at least 15-20 single cells from 5 individual brains were quantified in each of 2 independent experiments. Student’s t test was used to test for significant difference (p < 0.05) vs. control groups in most experiments. One-way ANOVA was performed to determine significant difference (p < 0.05) for rhythmic expression levels.

### 8. In situ hybridization

We used a protocol similar to ^73^ using HCR probes designed and synthesized by Molecular Instruments. Probes recognized the single *Pdf* exon, the first intron of *GlyP* or the second intron of *toy.* Larval brains were dissected in PBS and fixed in 4% PFA for 20 minutes and then washed in PBST (0.1% Tween). Brains were dehydrated with a MeOH series of increasing concentrations on ice and stored in 100% MeOH at -20°C. The following steps were then performed on ice: Brains were rehydrated in a MeOH series of decreasing concentrations and washed twice in PBST before permeabilizing in 5% acetic acid for 5 minutes and washing in PBST. Brains were refixed in 4% PFA for 20 minutes, and then washed in PBST, followed by a 1:1 mix of PBST and 5x SSCT, and then 2 washes in 5x SSCT. Brains were then incubated in Hybridization solution (Molecular Instruments) for 5 mins at room temperature and then in fresh hybridization solution for 30 mins at 37°C. 0.4 μL of each probe was added to brains in 100 μL hybridization solution overnight at 37°C.

Hybridization solution was then removed from samples which were washed 5 times in Wash buffer (Molecular Instruments) at 37°C, and then twice at room temperature in 5X SSCT. Brains were then equilibrated in Amplification buffer (Molecular Instruments) for 5 minutes.

Amplification probes were prepared according to the manufacturer’s instructions with 1.5 μl of the relevant amplification probes added to brains in 50 μl of Amplification buffer. Samples were then left overnight. Samples were washed in 5x SSCT at room temperature the next day. Brains were mounted in SlowFade Gold and imaged on a Leica SP8 confocal microscope.

## Acknowledgements

We are indebted to numerous stock centers for sharing flies, antibodies and plasmids: the HHMI Janelia Farm Research Campus, the Vienna Drosophila Resource Center (VDRC), the Bloomington *Drosophila* Stock Center (supported by NIH P40OD018537), the Developmental Studies Hybridoma Bank (created by the NICHD and maintained at The University of Iowa), the TRiP stock center at Harvard Medical School (supported by NIGMS, NIH) and FlyORF ^68^. We thank Claude Desplan for Toy antibodies, Gerry Rubin’s lab for plasmids, Jia Ling for advice on basal promoters, Jens Rister for advice on generating mutant reporter genes, Matthieu Cavey for advice on confocal microscopy and Chris Hackley for advice on cloning. We thank Esteban Mazzoni, Tom Carew, Adam Carter, Claude Desplan, Erich Jarvis and Niels Ringstad for advice and support and Claude Desplan, Simon Kidd, Alison Ehrlich and Kyla Hamling for comments on the manuscript. We are indebted to Dogukan Mizrak and Marc Ruben for deciding to profile LN_v_s in altered electrical states and to Subhabrata Sanyal for sharing his activity reporter genes many years ago. This investigation was conducted in facilities constructed with support from the NIH National Center for Research Resources. Imaging was performed at the NYU Center for Genomics & Systems Biology. ZZ was partly supported by the MacCracken Program and a James Arthur Dissertation Fellowship from NYU’s Graduate School of Arts and Science. Work in the authors’ laboratories was supported by BSF grant 2015506 (SK and JB) and NIH grants GM125859 (SK) and GM136363 (JB).

